# Defining the prototypical DNA replication fork trap in bacteria

**DOI:** 10.1101/2021.07.20.453168

**Authors:** Casey J. Toft, Morgane J. J. Moreau, Jiri Perutka, Savitri Mandapati, Peter Enyeart, Alanna E. Sorenson, Andrew D. Ellington, Patrick M. Schaeffer

## Abstract

In *Escherichia coli*, DNA replication termination is orchestrated by two clusters of *Ter* sites forming a DNA replication fork trap when bound by Tus proteins. The formation of a ‘locked’ Tus-*Ter* complex is essential for halting incoming DNA replication forks. However, the absence of replication fork arrest at some *Ter* sites raised questions about their significance. In this study, we examined the genome-wide distribution of Tus and found that only the six innermost *Ter* sites (*TerA-E* and *G*) were significantly bound by Tus. We also found that a single ectopic insertion of *TerB* in its non-permissive orientation could not be achieved, advocating against a need for ‘back-up’ *Ter* sites. Finally, examination of the genomes of a variety of Enterobacterales revealed a new replication fork trap architecture exclusively found outside the Enterobacteriaceae family. Taken together, our data enabled the delineation of a narrow prototypical Tus-dependent DNA replication fork trap consisting of only two *Ter* sites.

**Graphical Abstract:** 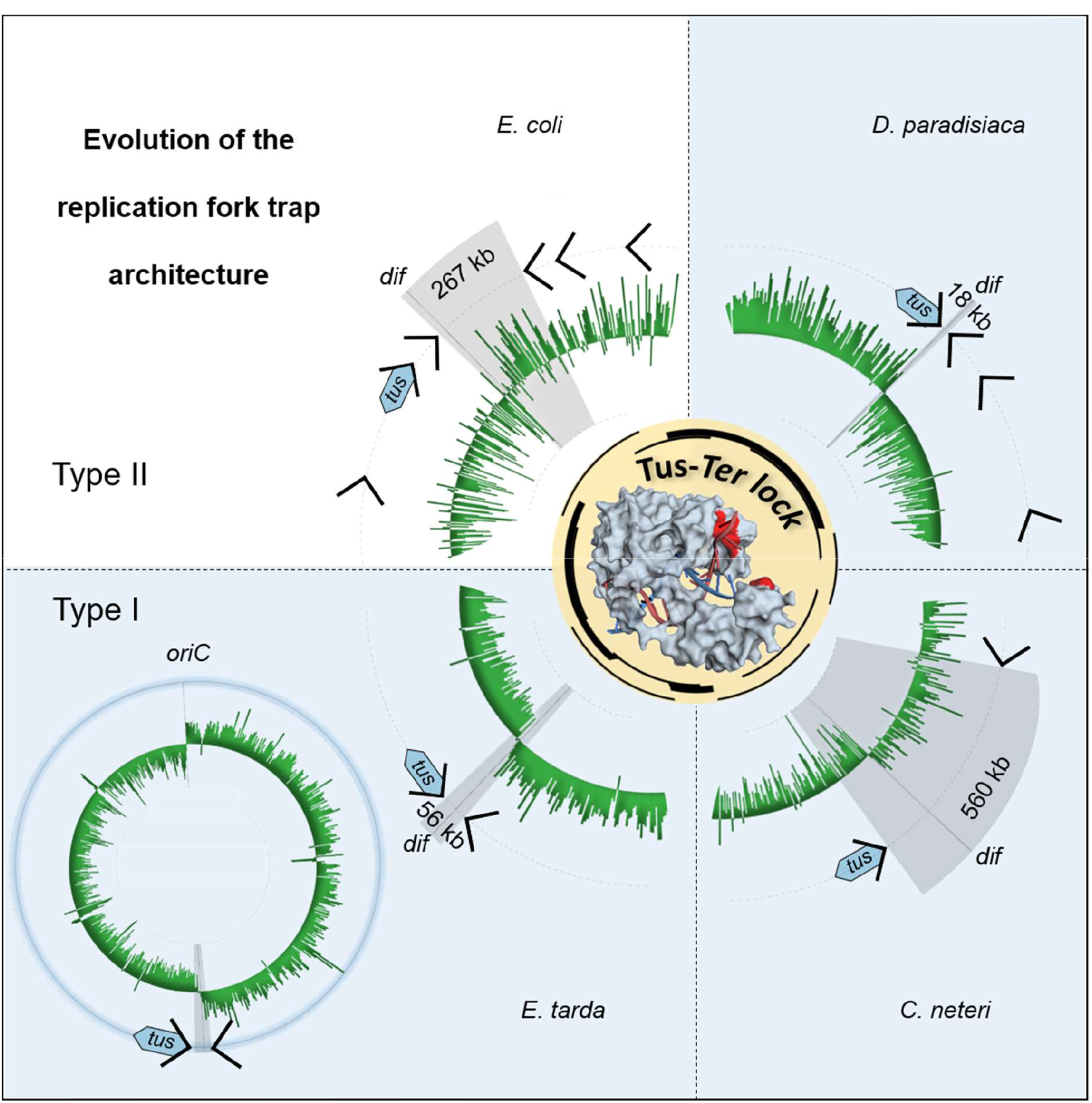

## INTRODUCTION

DNA replication termination in bacteria that utilise a DNA replication fork trap system within the chromosomal terminus region have been intensely scrutinized (1). This is exemplified in *E. coli* by the presence of a cluster of five similar but distinct 23-bp *Ter* DNA sequences on each chromosomal arm, which have anti-helicase activity when they are bound by the replication termination protein Tus (1–3). The complexity of the *E. coli* replication fork trap with respect to multiplicity and wide distribution of *Ter* sites around the chromosome is puzzling. One cluster consisting of *TerB*, *C*, *F*, *G* and *J* arrests the clockwise moving replication fork and the other oppositely-oriented cluster consisting of *TerA*, *D*, *E*, *I*, *H* arrests the anti-clockwise moving replication fork (Figure 1A). Until recently, the notions that Tus could bind to all ten of these slightly different *Ter* DNA sequences (*TerA-J*) (Figure 1B), and that these sequences all have a significant role in replication termination, have remained mostly unchallenged despite their individual binding properties for Tus being significantly different (4). Each *Ter* cluster consists of three high affinity, one moderate-to-low affinity and one non-lock forming *Ter* site (Figure 1A) (5, 6). Four additional *Ter*-like sequences (*TerK*, *L*, *Y* and *Z*) can be found in the *E. coli* chromosome, one within the previously identified termination region and the other three being on the left part of the chromosome, but these were dismissed as pseudo-*Ter* sites (7). Binding of Tus to the pseudo-*Ter* sites is likely to be insignificant based on their sequences (6) and fork arrest efficiency (7).

**Figure 1:**
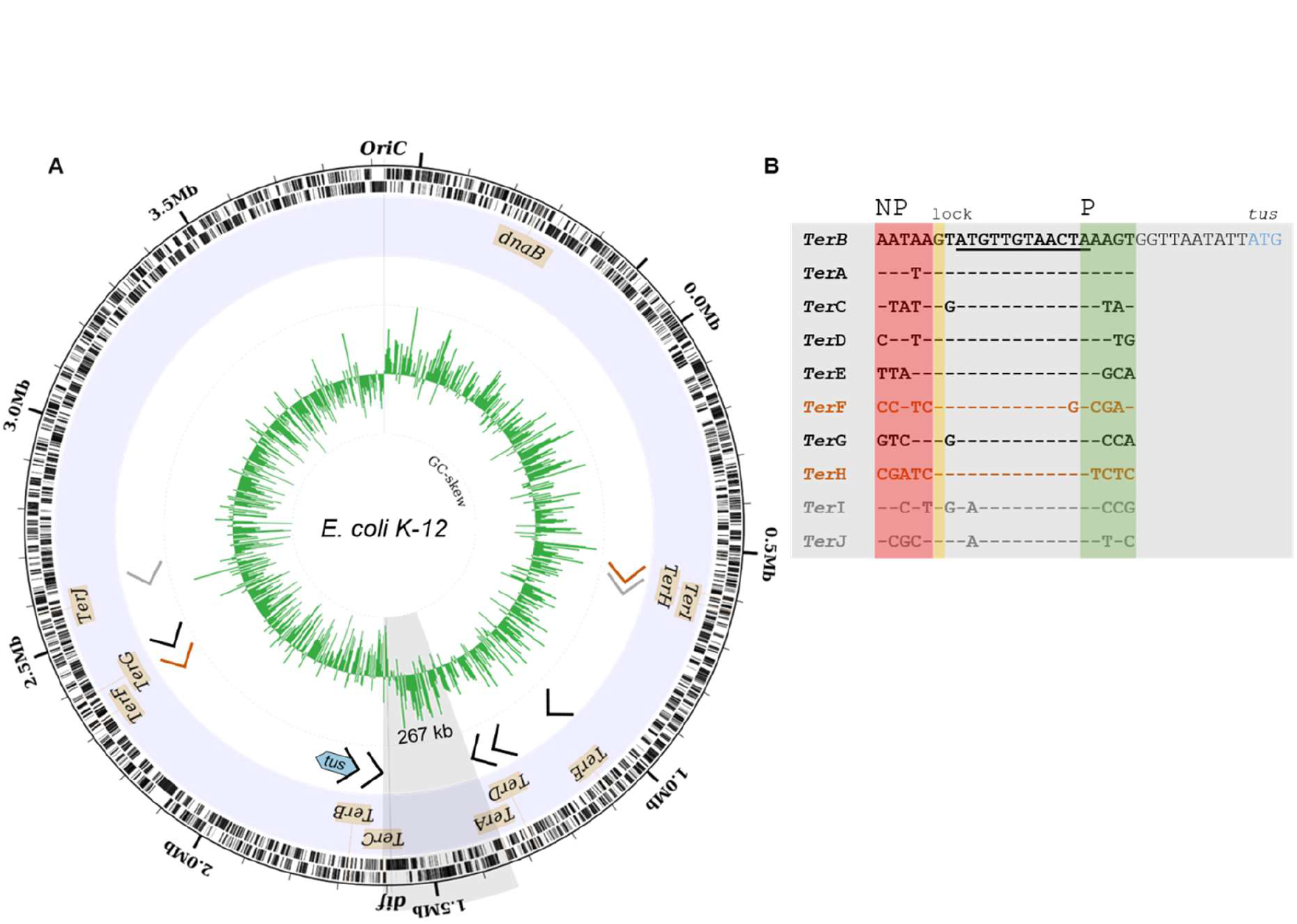
Chromosomal distribution and sequences of *Ter* sites in *E. coli*. (A) Circular representation of *E. coli* K12 MG1655. Illustrated from the outside to the centre of the circle: labelled forward and reverse genes; location of the ten primary *Ter*-sites (*TerA*-*J*) and their strand orientation; the currently accepted replication termination fork trap involving high affinity (black), moderate affinity (grey) and non-lock forming *Ter* sites (orange); GC-skew over a 5000 bp window showing a switch in polarity at the replication origin and close to *TerC* and the *dif* site. (B) *Ter* site sequences with the G(6) base complementary to C(6) highlighted in yellow and the strictly conserved 12 bp core sequence (underlined). *TerB* is located 11 bp upstream of the start codon (ATG) of the *tus* gene. NP: non-permissive face (red), P: permissive face (green).

The unique mechanism of polar DNA replication fork arrest observed in *E. coli* is due to the unusual binding mode of Tus to *Ter* and the unwinding action of the DnaB helicase at the non-permissive face of the Tus-*Ter* complex (1,7–12). Although a specific protein-protein interaction between the DnaB helicase and Tus had initially been proposed to have a pivotal role in polar fork arrest (8, 13), several studies have shown that this interaction is not necessary for polar fork arrest at the non-permissive face of the Tus-*Ter* complex (6,10,14–16). Tus precisely and tightly binds onto a *Ter* site, bending the DNA to prepare the molecular mouse trap that will be triggered by the 5’-3’ translocation and DNA unwinding action of DnaB helicase on the lagging strand moving towards the non-permissive face of the Tus-*Ter* complex (6,10,17). The progressive separation of DNA strands at the non-permissive face of the Tus-*Ter* complex ultimately breaks the GC(6) base pair (Figure 1B) in the *Ter* core sequence leading to the precise docking of the freed C(6) into a cytosine-specific binding pocket on the surface of Tus. The formation of the locked Tus-*Ter* conformation (TT-lock) slows the dissociation of Tus considerably and is believed to inhibit further DnaB helicase translocation (6,9,10). By using a systematic and comparative approach examining the kinetic and equilibrium parameters of all ten Tus-*Ter* and locked complexes, the binding of Tus to *Ter* and formation of the locked complex could be dissected in further detail (6,15,18). The study proposed a sequential three-step model for fork arrest including initial non-specific ‘sliding’ of Tus on DNA mediated by weak cooperative electrostatic interactions, followed by proper ratchet-like docking of Tus onto *Ter* upon correct alignment of specific nucleotide-amino acid contacts, and finally the DnaB-induced Tus-*Ter*-lock via binding of C(6) to the cytosine binding-pocket of Tus (6). The same study provided a new classification of *Ter* sites based on their kinetic and affinity parameters as well as their capacity to form a locked complex, and challenged the status quo by rejecting *TerF* and *TerH* as functional *Ter* sequences arguing that they could not induce polar fork arrest due to their inability to form a locked complex. A substitution of the canonical T to G at position 5 in the core sequence (Figure 1B) was proposed to be the cause for this loss of function (6). In addition, Coskun-Ari and Hill had previously observed a significant loss of replication fork arrest activity when *TerB* was mutated at TA(9) to CG or GC (19). *TerI* and *TerJ* are the only *Ter* sites where the TA(9) is replaced by an AT, which taken together with the relatively fast Tus dissociation from these locked Tus-*Ter* complexes (6) also raises doubts about their actual role in replication termination under natural Tus abundance conditions.

Using two-dimensional gel analysis of replication intermediates at *Ter* sites under natural conditions, Duggin and Bell showed that *TerC* is the most frequently used, with significant fork pausing also observed at *TerA* and *TerB* (7). Tus over-expression was required to observe intermediates at some of the other *Ter* sites (7). The sites varied significantly in their capacity to arrest replication forks and this was later correlated to their respective affinity for Tus (high, moderate or weak) and their ability to form a TT-lock (5,6,18). Most puzzling, however, was the observation that some pausing occurred at the outer *TerH.* Indeed, to be arrested at *TerH*, the replication fork has to break through the strong *TerE* and moderate *TerI* but no fork pausing was observed at *TerE* and little at *TerI*. Duggin and Bell also showed that pausing was abolished at *TerC* in a *tus* null strain, but they did not verify if the pausing observed at the outermost *TerH-I* sites was also strictly due to Tus binding (7). The low probability of the anti-clockwise fork reaching *TerH*, the absence of pausing at the strong *TerE* (7) and the non-TT-lock forming characteristic of *TerH* (6), suggest that the pausing observed at *TerH* could either be due to a clockwise moving fork at the permissive face of *TerH* or to recombination events (20–22).

The presence of the distal *Ter* sites and their involvement in DNA replication termination remains unclear. Forks most frequently meet at *TerC* and to some extent at *TerA* and *B* (7). Assuming the two forks progress at equivalent rates, forks are more likely to meet at *TerC* than at *TerA* since *TerC* is almost perfectly located directly opposite to *oriC* whereas the anti-clockwise moving fork must travel an additional ∼259 kb to encounter the non-permissive face of the Tus-*TerA* complex. Despite the stability of the locked Tus-*TerC* complex (6) in over-expressed Tus conditions, significant pausing still occurs at *TerB* and to some extent at *TerG* (7). One possible explanation for pausing at *TerB* is that in some cases, the ratchet-lock mechanism (6) fails to form and the next site serves as a backup for DNA replication arrest. In support of this, single molecule DNA replication assays suggest that a replication fork approaching a non-permissive *TerB* will fail to be arrested 52% of the time as a result of an inefficient Tus-*Ter*-lock mechanism (16). The authors proposed that lock formation is dependent on transient fork stoppage by an Arg198 interaction that buys time for C(6) to dock into its binding pocket (16). Hence the critical need for back-up *Ter* sites throughout the terminus of the genome is for replication forks that have breached the innermost *TerA* and *TerC* sites.

So far the large number of binding, structural and single molecule studies designed to thoroughly examine the Tus-*Ter* complexes (2,5,6,15,16,23), as well as fluorescence imaging aiming at examining the progression and pausing of replication forks at natural replication barriers in live bacteria (24), have failed to provide a clear explanation for the need of such a large replication fork trap in *E. coli*. In fact, the most significant knowledge gap on Tus-*Ter* replication fork traps, i.e. the binding distribution of Tus to individual *Ter* sites in replicating bacteria, has not been addressed. As such, the function of distal *Ter* sites and their biological relevance remains unclear and calls into question as to whether or not Tus proteins (25, 26) really bind these *Ter* sites, and if yes, then to what extent?

Here we examined the impact of inserting a selection of *Ter* sites (*TerB*, *H* and *J*) into a safe chromosomal locus in both permissive and non-permissive orientations using TargeTron technology, as well as the genome-wide distribution of Tus using ChIP-Seq and ChIP-qPCR to identify the functional *Ter* sites capable of halting replication forks in *E. coli*. We also characterised the fork trap architecture in closely, moderately, as well as distantly related bacteria harbouring the *tus* gene. Taken together, our data enabled the delineation of a prototypical Tus-dependent DNA replication fork trap.

## MATERIALS AND METHODS

### Materials and other resources

See Supplementary Data.

### Plasmids and protein expression

The vector encoding His_6_-Tus-GFP (pPMS1259) has been described previously (27–30). *E. coli* KRX (Promega) which is a K12 derivative was used to express His_6_-Tus-GFP. Reference His_6_-Tus-GFP proteins were expressed and affinity purified as previously described (29), and stored in 50 mM sodium phosphate (pH 7.8) and 10% glycerol (w/v).

### Protein expression and crosslinking

The ChIP protocol was derived from previous work by Regev *et al.* and Ishikawa *et al.* (31, 32). Competent *E. coli* KRX bacteria were transformed with pPMS1259, plated onto LB plates supplemented with ampicillin (100 µg/ml) and grown overnight at 37°C. Colonies were resuspended and diluted to an OD_600_ of 0.1 in 12 ml of LB broth supplemented with ampicillin (100 µg/ml). Cultures were grown for 45 minutes at 37°C before inducing moderate levels of His_6_-Tus-GFP with 0.02 % Rhamnose (w/v final culture concentration). Cultures were incubated for another 2 hours at 37°C, followed by 2 hours at 16°C. Culture aliquots (9 ml) were cooled on ice for 30 minutes to which 250 µl of a formaldehyde solution (36% w/v) were added (final concentration of 1%). The bacterial suspensions were then placed at room temperature for 20 minutes. Glycine powder was added the bacterial suspensions (0.5 M final concentration) for 5 minutes at room temperature followed by 5 minutes on ice. The bacterial suspensions were centrifuged 5 minutes at 800 *g* and 4°C and washed twice with 4 ml and 10 ml of cold TCS buffer (50 mM Tris (pH 7.5), 150 mM NaCl and 2 mM KCl). The supernatants were discarded and the bacteria pellets were stored at -80°C until required. KRX bacteria (without plasmid) were subjected to the same protocol and used as control.

### Detection and quantitation of GFP-tagged protein expression

Culture aliquots were taken prior to crosslinking, centrifuged at 1,000 g for 1 minute and the pellets were resuspended in 2X Laemmli buffer at a concentration of 7.8 x 10^9^ bacteria.ml^-1^. The suspensions were heated for 10 minutes at 90°C and 5 µl (corresponding to total proteins of 3.95 x 10^7^ cells) were separated in a 10 % SDS-polyacrylamide gel alongside 0.5 µg of reference His_6_-Tus-GFP protein standard for Western blot analysis. Chicken anti-GFP IgY (Abcam ab92456) and HRP-conjugated goat anti-IgY (Jackson 103-035-155) were revealed with SIGMAFAST™ 3,3′-Diaminobenzidine tablets. Protein bands were quantified using imageJ (http://rsbweb.nih.gov/ij/) and intracellular concentrations were estimated based on the intensity of bands of known protein concentration and using cell parameters determined by Volkmer and Heinemann for cell volume (4.4 fL) and cell concentration at a given OD_600_ in LB (7.8×10^8^ cells.ml^-1^.OD^-1^) (33).

### Chromosome immunoprecipitation

Bacteria pellets were resuspended in lysis buffer (10 mM Tris (pH 8), 20 % sucrose, 50 mM NaCl, 10 mM EDTA, 1 mg/ml lsozyme and 10 µg/ml RNase) in 1/10 of initial culture volume (adjusted between replicates to reach same suspension concentration). Following a 30-minute incubation period at 37°C, the lysates were diluted 5 times in IP buffer (50 mM HEPES-KOH (pH 7.5), 150 mM NaCl, 1 mM EDTA) and passed three times in a French press at 12,000 psi to ensure maximum and reproducible cell lysis and DNA shearing. The Tus-GFP lysates were heated for 10 minutes at 50°C to denature free Tus-GFP (6,28,29). Control KRX lysates were processed identically. All lysates were centrifuged at 30,000 *g* for 20 minutes at 4°C. Supernatants were used for immunoprecipitation and as input samples. For immunoprecipitation, 96-well MaxiSorp round bottom U96 Nunc plates were coated overnight at 4°C with 50 µl of 50 mM sodium phosphate (pH 7.5) and 10% glycerol buffer containing 0.5 µg of goat anti-GFP IgG (Abcam; Ab6673). Wells were washed once with 200 µl of TCS buffer prior to immunoprecipitation. After the wash step, 50 µl of lysate supernatant were added per well for 90 minutes at room temperature. Wells were washed three times with 200 µl of TCS buffer. Control immunoprecipitation experiments were performed in parallel without antibody pre-coating as background controls. Immunocaptured DNA was released by adding 50 µl of elution and de-crosslinking buffer (2 mM Tris (pH 7.5), 50 mM NaCl, 0.005 % tween and 300 µg/ml proteinase K) to each well for 1 hour at 37°C (output). Input samples were diluted 10,000 times in elution buffer (2 mM Tris (pH 7.5), 50 mM NaCl, 0.005 % Tween) and 50 µl were transferred to a tube containing proteinase K yielding a final concentration of 300 µg/ml to de-crosslink the input DNA for 1 hour at 37°C. Samples (inputs and outputs) were incubated 15 minutes at 95°C to denature proteinase K and residual crosslinked proteins. After 5 minutes incubation on ice, samples were centrifuged at 18,000 *g* for 5 minutes at 4°C and the supernatants were used for qPCR and Illumina sequencing.

### qPCR protocol

All qPCR reactions were performed as previously described (5). Oligonucleotides for amplification of *oriC* and *Ter* containing regions are listed in Additional Resources. Briefly, qPCR reactions contained 2 µl of input or output DNA sample, 8 µl of primer pairs (0.5 µM each) and 10 µl of SensiMix SYBR & fluorescein mastermix (Bioline). The protocol included a 10 minute step at 95°C followed by 40 cycles at 95°C for 10 s and 60°C for 15 s. Melt-curves were run for quality control. *Ct* values were obtained at a set threshold applied to all experiments. Standard curves were performed in triplicate with purified and serially diluted *Ter* and *oriC* amplicons in matching output buffer conditions. For each primer pair, the average slope value of three standard curves (n=3) was used to determine the primer specific amplification efficiency according to the following equation (34).

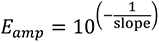

A melt-curve was performed to verify that the correct regions were amplified.

### qPCR analyses

For all qPCR experiments, *Ct* values were determined at the same threshold value. *Ct_(input)_* values were corrected for the dilution factor to give *_c_Ct_(input)_* according to the following equation:

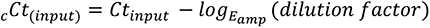

The immunoprecipitation efficiency of each specific target DNA region relative to a non-specific DNA region (*IP efficiency_(oriC)_*) was calculated according to the following equation:

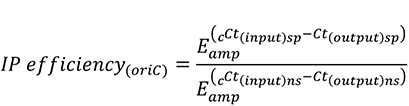

where *_c_Ct_(input)_* and *Ct_(output)_* are values obtained for each DNA target before (input) and after ChIP (output). Specific DNA target (i.e. *Ter* sites) and non-specific control DNA region are indicated with “sp” and “ns” subscripts respectively.

### Library preparation and Illumina sequencing

Input and output DNA samples were purified using Wizard PCR clean up and eluted in 110 µL water. Each library was prepared using the NEBNext Ultra DNA library preparation kit for Illumina. ChIP output samples (∼0.25 ng) were used for library preparation. The libraries were prepared according to manufacturer instructions. Briefly, 55.5 µL of DNA suspensions were end repaired. Due to the low DNA concentrations in the suspensions, the NEB adaptors were diluted 10 fold in water to 1.5 µM for ligation as recommended. The adaptors were cleaved using uracil excision. Size selection was not recommended for sample < 50 ng. DNA was then cleaned up using Sera-Mag beads (ratio of 1.4) and eluted in 28 µL of 0.1xTE. Index primers were added by PCR using 18 cycles. (13-15 cycles were recommended for 5 ng of input material, therefore 3 cycles were added to account for the 20 fold difference in input DNA). DNA quantification was performed using the Quantifluor dsDNA system (Promega). The samples were pooled in a single library, denatured and loaded for sequencing with an Illumina MiSeq desktop sequencer (50 bp single-end sequencing). Illumina read quality was assessed using FastQC (v.0.11.8) followed by removal of Illumina adapters and leading and tailing nucleotides with a Phred score ≤ 10 over a 6 bp window using Trimmomatic (v.0.36).

### DNA preparation for Nanopore long read sequencing

A flask containing 10 ml of LB media was inoculated with 100 μL of KRX *E. coli* overnight culture. The culture was incubated at 37°C and 150 RPM until log phase was reached (OD_600_ = 0.7) at which point chloramphenicol was added at a final concentration of 180 μg/mL to inhibit protein synthesis. The bacteria were centrifuged and resuspended in 3.5 mL lysis buffer (114 mM Tris-HCl (pH 8), 115 mM EDTA, 570 mM NaCl and 1% triton X-100). After addition of lysozyme (250 μL, 50 mg/mL), sodium dodecyl sulphate (500 μL, 10% w/v) and RNase A (4 μL, 100 mg/mL), the bacterial suspension was inverted gently 12 times and heated at 50°C for 30 min. Proteinase K (500 μL, 10 mg/mL) was added with repeated gentle mixing and heating steps. The suspension was then combined with 8 mL precipitation buffer (75% isopropanol, 2.5 mM ammonium acetate), inverted gently 15 times followed by centrifugation at 4,000 *g* at 4°C for 5 min. The supernatant was discarded. The soft DNA precipitate was transferred using a wide bore tip into a 1.5 mL tube, resuspended with 70% ethanol and stored at 4°C for 5 min, then centrifuged and the supernatant removed. The DNA pellet was air dried for 5 minutes then resuspended in 200 μL nuclease free water and heated at 50°C for 5 minutes with the tube cap open. The DNA was sheared using a syringe with a 20 gauge needle (3 times). DNA concentration and quality were assessed using Invitrogen Qubit 4 and agarose gel electrophoresis.

### Nanopore long read sequencing and genome assembly

A Nanopore sequencing library was prepared using the Rapid Sequencing protocol SQK-RAD004 (Oxford Nanopore). As recommended by the protocol, 7.5 µl of DNA suspension (400 ng) was added to the flow cell. Sequencing was performed on a FLO-MIN106 R9 MinION flow cell. Base-calling was processed using the pipeline implemented in MinKNOW software version 18.01.6 (Oxford Nanopore). In total, 1.17 GB (253x coverage) of sequence data was generated for *E. coli* strain KRX over ∼18 hours of sequencing achieved in two separate runs. Prior to assembly, all fastq files were combined and quality filtered by nanofilt version 2.5.0 (quality score ≥ 9). The remaining 225 985 reads had an average length of 3738 bp with the longest read of 179 912 bp and a total 146x coverage of the KRX *E.coli* genome. Oxford Nanopore adapters were trimmed using Porechop (version 0.2.3_seqan2.1.1) and assembled using Flye (version 2.6) with default settings for Oxford Nanopore data. Polishing was performed with Racon iteratively four times in combination with Pilon (version 1.23-1) using the entire Illumina ChIP-Seq data (i.e. KRX Input, WGS, negative control and ChIP DNA). The genome was annotated using Prokka (version 1.14.0) and evaluated using Quast (version 5.0.2) against the *E.coli* K12 genome (GenBank assembly accession: GCA_000005845.2).

### ChIP-Seq analysis

Following adapter trimming and quality control, each biological and technical replicate fastqc file of sequenced samples (i.e. Input, ChIP DNA and negative control) were individually aligned to the polished KRX genome using Bowtie2 (version 2.3.4.1). Samtools (version 1.9) was then used to organise each alignment file for visualisation on Interactive Genomics Viewer (IGV) to visually assess the data in terms of replicability between each replicate and for any outliers. The replicate data for Input, ChIP and negative control were pooled into three separate fastqc files and aligned to the KRX *E. coli* genome using the default settings of Bowtie2. A circular annotation of each pooled ChIP-Seq reads mapped to the KRX *E. coli* genome spanning a 23 bp window (size of an extended *Ter* site) was created as well as the GC-skew of the chromosome (5000 bp window) using Circleator (version 1.0.2). The individual positions and orientations of *Ter* sites were identified and verified using their specific 23 bp sequences and compared to the respective sites in *E. coli* (K12). The single base read counts were averaged over the 23 base *Ter* sequences for Input, ChIP and negative control DNA using genomeCoverageBed (Version: v2.26.0).

### Genome engineering of ectopic *Ter* sites

Targetrons (mobile group II introns) were designed to insert *Ter* sites in the safe insertion region SIR.5.6 (35), located in the right non-structured chromosome domain (Figure 1A) of *E. coli* BL21(*DE3*) (accession number AM946981). The SIR5.6 is located about 930 kbp downstream of *oriC* (right chromosome arm). *Ter*-targetrons (mobile group II introns carrying *Ter* sequences) insertion was performed as described previously (36), replacing *lox* sites with *TerB*, *TerH* or *TerI* sequences in permissive (P) or non-permissive (NP) orientations. Insertion of *TerB* in the non-permissive orientation was also attempted using the Lambda Red recombination system but no viable colonies could be obtained. The successful insertions of the other *Ter* sites were confirmed by colony PCR and verified by sequencing. *Ter* sites were inserted into SIR5.6 with an efficiency of 53/65 (81.5% - excluding integrations attempted for the insertion of *TerB* in the non-permissive orientation).

BL21(*DE3*) cells carrying ectopic *Ter* sites were grown in LB broth at 37°C and OD_600_ was measured every 5 minutes for 12 hours. The results were plotted as log_2_(OD_600_) versus time (minute). In order to select the linear region of the curve, each point was assigned a correlation coefficient R^2^ corresponding to the value of R^2^ for the line consisting of that point and the five points before and after. The variance was lower when the same time window was used for all three replicates so the resulting R^2^ values were averaged for all three replicates at each time point. The longest stretch in which all these averaged R^2^ values were equal to or greater than 0.99 was taken as the linear range. The slope of the least-squares linear fit of the log_2_(OD_600_) curve of each replicate in that time range was then taken as the growth rate and the doubling time was calculated as 1/growth rate.

### Fork trap characterisation in Enterobacterales

Bacterial species containing a *tus* gene ortholog were identified using InterPro entry IPR008865 including only the entries for Enterobacterales (2518 protein sequences in total). Several Tus protein sequences from a selection of *Pseudoalteromonas* species were also included as outgroup. Ten rounds of an alignment for all sequences were generated using MUSCLE software with default settings (37). The tree was constructed using the matrix of aligned sequences in RAxML v8.2.0 (38) which performed an ML phylogenetic analysis with 100 independent repetitions using the PROTGAMMALGI molecular evolution model in combination with an independent rapid bootstrap algorithm (--AutoMRE) to establish support for each node. The consensus tree produced by RAxML was visualised and edited in iTol (39). The sequences that branched off earlier than the outgroup were removed from the final tree as these were most likely incorrectly assigned taxa or sequences that are most likely not Tus protein sequences. For clarity, clades from *Escherichia coli*, *Salmonella* and *Klebsiella* were collapsed due to the large number of sequences.

A random selection of species from different families were chosen including *Dickeya paradisiaca* (strain Ech703), *Edwardsiella tarda* (strain EIB202), *Yersinia pestis* (Microtus str. 91001), *Xenorhabdus nematophila* (strain ATCC 19061), *Proteus mirabilis* (strain HN2p), *Cedecea neteri* (strain ND14a) and *Salmonella typhimurium* (strain LT2). Genome assemblies that were preliminary were excluded from fork trap analysis. Upon identification of a *tus* gene otholog, the adjacent *Ter* site was identified within its 50 bp 5’ UTR by aligning the 23 bp *E. coli TerB* sequence. For each selected species, a BLAST search was carried out using the adjacent *Ter* sequence to locate further *Ter* sites within their genome. Sequences were verified by inspecting each BLAST to ensure it contained the locking C(6) followed by the conserved 12 bp core spanning from A(8) to A(19) (Figure 1B). A circular annotation of each genome to display the architecture of the fork trap was generated alongside the GC-skew of the chromosome (5000 bp window) using Circleator (version 1.0.2).

### Quantification and statistical analyses

Statistics and number of biological and technical repeats are indicated in the relevant figure legends, tables and methods. Statistical analyses were performed using Graphpad Prism 7. Data are expressed as mean values ± SD, ± SE or ranges.

## RESULTS

### Ectopic insertion of *TerB*, *TerH* and *TerJ* sites

The role of some of the distal *Ter* sites in replication fork arrest is questionable in light of their location, Tus binding affinity and dissociation kinetics (6). *TerF* has recently been dismissed as a pseudo-*Ter* site with no possible role in replication fork arrest. While *TerH* cannot form a locked complex, the locked Tus-*TerJ* complex has a dissociation half-life (t_1/2_) of 332 s that matches the non-locked Tus-*TerB* t_1/2_=315 s at 250 mM KCl (6). These findings prompted us to examine the capacity of the most distal *Ter* sites (i.e. *TerH* and *J*) to halt DNA replication forks. For this, *TerH* (moderate affinity, non-TT-lock forming sequence, t_1/2_=59 s), *TerJ* (weakest moderate affinity, weak TT-lock forming sequence, t_1/2_=332 s) and the strong TT-lock forming *TerB* (t_1/2_=4367 s) were inserted in the right chromosome arm of *RecA^+^ E. coli* strain BL21(*DE3*), 930 kbp downstream of *oriC* (right arm, SIR5.6) in both permissive (P) or non-permissive (NP) orientations using a TargeTron strategy (36). While ectopic *Ter* site insertions and fork trap inversions have been studied previously (40–43), the TargeTron technique guarantees that a 23 bp *Ter* within a short intron sequence is incorporated with minimal genomic variations. We hypothesised that *Ter* insertions resulting in weak to moderate replication fork pausing would yield a measurable effect on bacterial growth rate while *Ter* insertions yielding efficient ectopic fork arrest should be unviable. As such, ectopic insertion of *TerB* should be fully viable in permissive orientation and unviable in non-permissive orientation if the TT-lock is unbreachable.

Growth rates were determined for viable *E. coli* cells with successful ectopic *Ter* sites insertions confirmed by sequencing (Table 1). All *Ter* sites, except *TerB* in non-permissive orientation, could be inserted in either permissive or non-permissive orientation into SIR5.6. *Ter* sites were inserted with an efficiency of 53/65 (81.5% - excluding integrations attempted for the insertion of *TerB* in the non-permissive orientation). It is important to note that *TerB* in non-permissive orientation could not be inserted using either a TargeTron or the Lambda Red recombination system (44). All viable strains reached the same plateau at the same time as the control strain (SI Figure 1) suggesting that *TerB* in permissive orientation as well as *TerH* and *TerJ* in either orientation do not impact replication forks or chromosomal segregation which is in partial agreement with previous genomic region inversion data (43). Furthermore, no significant difference in bacterial growth rates or delays were observed between these and the control strain. We conclude that the site-specific insertion of an ectopic *TerB* in non-permissive orientation in a strain carrying the wild type *tus* gene is unviable as a result of efficient replication fork arrest 930 kbp downstream to *oriC.* We presume that the resulting fork stalling or reversal induced by *TerB* in non-permissive orientation cannot be resolved, even in a *RecA^+^* strain with wild-type homologous recombination function (45).

**Table 1:** Effect of ectopic Ter sites on the growth rate of *E. coli* BL21(DE3)

| Strain | Growth rate | Doubling time |
| --- | --- | --- |
| BL21(DE3) | $\text{Log}_2(A_{600})/\text{min}$ (SD) | $T_D$ in min (SE) |
| *TerB* (P) | 0.039 (0.0012) | 25.53 (0.448) |
| *TerH* (NP) | 0.041 (0.0002) | 24.13 (0.087)* |
| *TerH*(P) | 0.042 (0.0017) | 23.69 (0.564) |
| *TerJ* (P) | 0.044 (0.0011) | 22.70 (0.327) |
| *TerJ* (NP) | 0.042 (0.0008) | 23.70 (0.259) |
| Control | 0.039 (0.0012) | 25.88 (0.469) |

*TerB*, *TerH* and *TerJ* were inserted ~ 930 kpb downstream of *oriC* in the permissive (P) or non-permissive (NP) orientation. A culture of wild type BL21(DE3) was grown as a control. Growth rates were determined from the slopes of the linear regressions performed between 100 and 210 minutes. Doubling times ( $T_D$ ) were calculated as $1/\text{growth rates}$ ( $n=3$ , except for *TerH* (NP), $n=2$ ). Standard deviations (SD) are shown. \* One outlier was omitted.

Our data support the notion that *TerH* and *J* do not arrest nor pause replication forks and as such we propose to reclassify them as pseudo-*Ter* sequences. Furthermore, in light of our and previous data (6,7,19), *TerI,* which forms a faster dissociating TT-lock complex (t_1/2_=196 s) than *TerJ* (t_1/2_=332 s) at 250 mM KCl (6, 18), can also reasonably be dismissed as a pseudo-*Ter* site.

### Chromosomal binding of Tus

The genome wide distribution of Tus was examined by using chromosome immunoprecipitation (ChIP)-Seq and ChIP-qPCR (SI Figure 2) to identify the *Ter* sites that are bound during active DNA replication. Due to the low natural abundance of Tus and the unavailability of Tus-specific antibodies, the chromosomal distribution of GFP-tagged Tus (Tus-GFP) was examined in exponentially growing *E. coli* (KRX). Tus autoregulates its expression via binding to *TerB* located within the promoter region of the chromosomal *tus* gene (25, 26). As such, we hypothesized that in the presence of excess Tus-GFP, the transcription of the *tus* gene would be downregulated further. This would have the effect of reducing *in vivo* Tus levels allowing excess Tus-GFP to efficiently compete for *Ter* sites. Due to the unique base sequences flanking each *Ter* site, ChIP samples could be sequenced using 50 bp Illumina reads, thereby ensuring that the reads containing the full or partial 23 bp *Ter* sequences could be accurately mapped to the genome. Input and immunoprecipitated DNA samples were sequenced and the reads mapped back to our KRX genome assembly to generate a high-resolution genome-wide distribution map of Tus-GFP (Figure 2).

**Figure 2:**
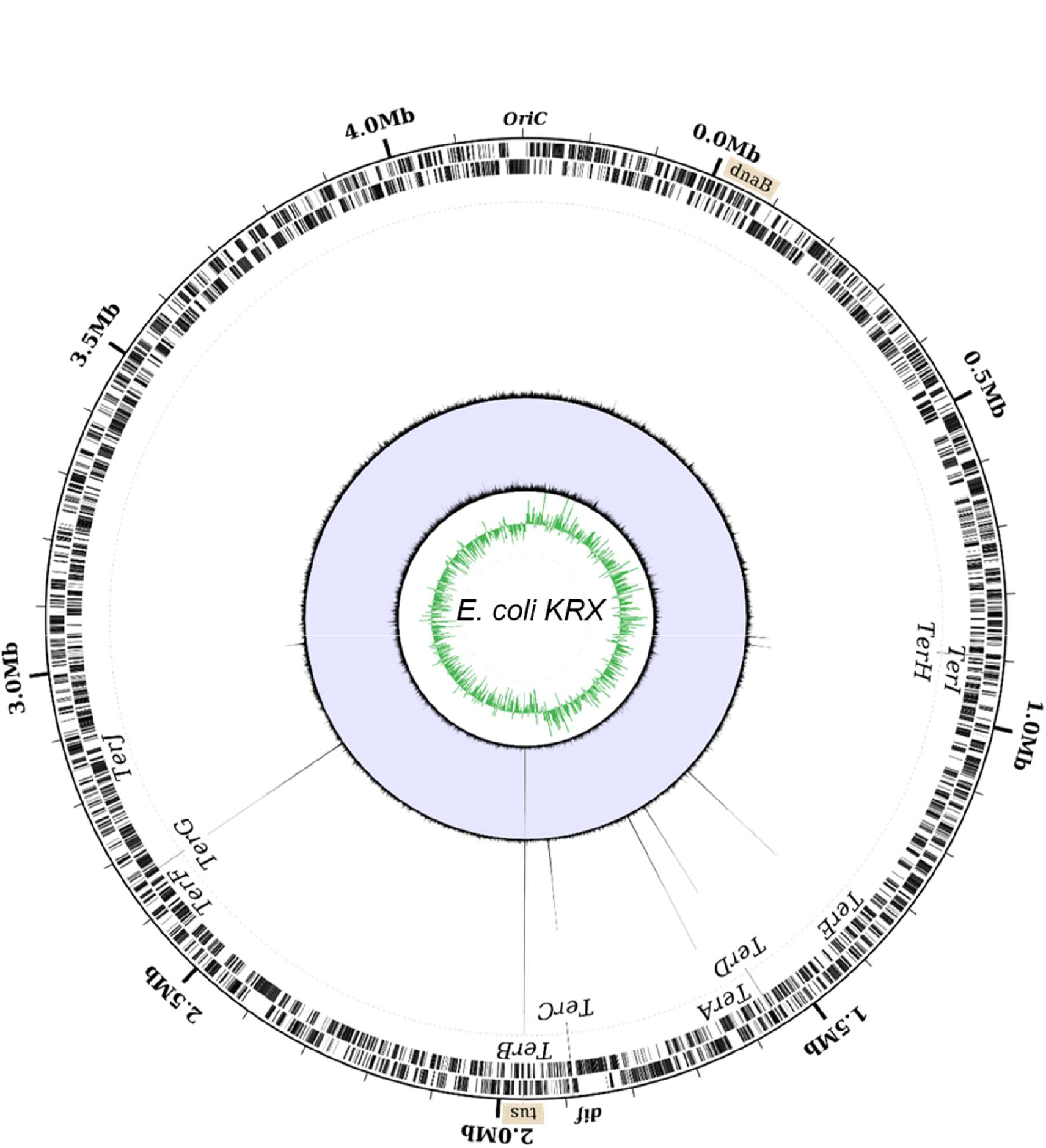
Circular representation of *E. coli* KRX chromosome with mapped ChIP-Seq coverage. From the outside to the centre of the circle: labelled forward and reverse genes; genomic location of sites and genes involved in DNA replication termination; combined ChIP-Seq read coverage (max = 430 reads at *TerB*), Input DNA read coverage (max = 230 reads at the *tus* gene), GC-skew over a 5,000 bp moving window. The GC-skew switches polarity at the replication origin and terminus.

A large peak at the *tus* locus was clearly visible in the input DNA sample which corresponded to the plasmid-encoded *tus* sequence counts. The coverage at *tus* in the input DNA indicated a plasmid copy number per chromosome of ∼ 43. ChIP-Seq peaks were immediately apparent without the need for a peak identification workflow. Six large peaks were visible in the immunoprecipitated coverage plot, corresponding to the binding of Tus-GFP to individual *Ter* sites. We inspected the read coverage in the 10 bp region between the chromosomal *tus* and *TerB* loci to ensure that the reads originating from plasmid-encoded *tus* did not bias the read count at *TerB* (SI Figure 3). The base coverage at *TerB* is equivalent to the average chromosomal reads in the input DNA demonstrating that our method does not lead to a coverage bias in the immunoprecipitated DNA data. The average coverage values ranged from ∼5 at *oriC* to ∼1 in the terminus region indicating that at least three replication forks were progressing on each chromosome arm towards the terminus region. Our ChIP-seq data revealed that out of the 10 primary *Ter* sites, only the 6 high-affinity *TerA-E and G* sequences (6) are significantly bound by excess Tus-GFP (Figure 2 and SI Figure 4).

Surprisingly, despite being the major termination site (7) *TerC* was one of the least bound in this group with an average 269x read coverage compared to 430x coverage at *TerB*. The coverage at *Ter*G (410x) was similar to *TerB* suggesting that this site is almost certainly bound at normal bacterial Tus concentrations. Given the strong Tus binding and lock-forming ability of *TerG* (6), our data suggest that the absence of paused fork intermediates in the fork arrest assay measured by Duggin and Bell (7), is a result of the replication fork not reaching this *Ter* site. As anticipated, no binding was observed at the pseudo-*TerF* (5-7,29). Out of the three moderate *Ter* sequences, *TerJ* in its locked complex with Tus is the most stable with respect to t_1/2_ (6) and fork arrest activity (7) yet no peak was observed strongly supporting our ectopic insertion data and that the latter is also a pseudo-*Ter* site in natural conditions. *TerH* and *TerI* sites have similar coverages (57x and 48x respectively) corresponding to only 11-13% of the coverage at *TerB*, despite the bacterial Tus-GFP concentration being 1,000-fold higher than the normal endogenous levels of Tus, suggesting these sites would be mostly unbound at normal cellular Tus concentration. Taken together with previous affinity data, our ChIP-Seq and ectopic insertion findings support the notion that *TerH, I* and *J* do not have a role in replication fork arrest. Our ChIP-Seq dataset was confirmed by ChIP-qPCR (SI Figure 5) and allowed delineation of a refined minimal replication fork trap within *E. coli* comprising two clusters of three *Ter* sites: (a) *TerB*, *C* and *G* that can arrest a clockwise moving replication fork and (b) *TerA*, *D* and *E* that can arrest an anticlockwise moving replication fork (Figure 2).

### GC-skew relative to Termination site usage in *E. coli*

Although the GC-skew is a well-recognised tool to identify the origin of replication in many circular prokaryotic chromosomes (46–48), the feasibility of utilising the GC-skew to predict the terminus has been debated amongst researchers due to the terminus shift point being closer to the chromosome dimer resolution site (*dif*) than to the *Ter* sites in some studied species (49). However, the GC-skew has recently been shown to coincide with replication fork arrest by Tus at *Ter* sites and is not influenced by *dif* (*50, 51*). We hypothesized that the GC-skew should correlate with the frequency of fork arrest activity at specified *Ter* sites. In other words, the GC-skew is representative of the average of the ensemble of replication forks collision loci at functional *Ter* sites. In this scenario, the inflection point should occur at the historical positional average between the *Ter* sites where termination occurs. Duggin and Bell showed that only *TerA, B* and *C* have significant replication arrest activity (0.19 %, 0.14 % and 0.85 % respectively) in natural Tus conditions (7). We tested this scenario and found that the expected average position of replication termination (based on the positional and fractional distribution of replication fork arrest activity) almost coincided with the GC-skew inflection point, i.e. only 7.5 kb from the calculated inflection point derived from a sliding 1,000 bp cumulative GC-skew (SI Figure 6). It is important to note that the *dif* site is located 8 kbp from the terminal GC-skew switch point on the other chromosomal arm. We tested additional scenarios but none produced a better correlation. Taken together with previously published data (50, 51), the GC-skew of *E. coli* supports the involvement of *TerA*, *B* and *C* in replication fork arrest and provides an invaluable tool to further our understanding of replication termination in other species.

### A narrow fork trap dyad in *Edwardsiella tarda*

While the function of *TerA*, *B* and *C* and their replication fork arrest activity in *E. coli* is clear, the need of *TerE, D* and *G* is not, despite their high affinity for Tus and ability to form a TT-lock (6). While trying to gain further insight into these seemingly redundant *Ter* sites, we examined the replication fork trap architecture in closely, moderately as well as distantly related bacteria harbouring a *tus* gene (Figure 3A-B). A recent phylogenetic analysis of Tus homologs in bacteria identified resident *tus* genes within the chromosomes of most Enterobacterales (52). Using a streamlined approach we characterised the replication fork traps in several of these species (SI Figure 7 & SI Table 1). Our approach used a refined definition of what constitutes a *Ter* site: (a) a 23 bp *Ter* sequence is always located within 50 bp upstream of the *tus* gene; (b) it must contain a GC base-pair at position 6 followed by the conserved 12 bp core spanning from AT(8) to AT(19) (Figure 1B) and (c) it cannot contain a G at position 5 next to C(6) to ensure unhindered formation of the locked complex with Tus. Following identification of the vicinal *Ter* sequences upstream of *tus* genes in our selected bacterial genomes, BLAST searches were performed to identify other *Ter* sites within the genomes as well as their replication fork blocking orientations. The stringency of our approach was evaluated with *E. coli* K12, identifying all primary *Ter* sites as well as the pseudo-*TerH*-*J* but excluding the pseudo-*TerF, K*, *L*, *Y* and *Z*.

**Figure 3:**
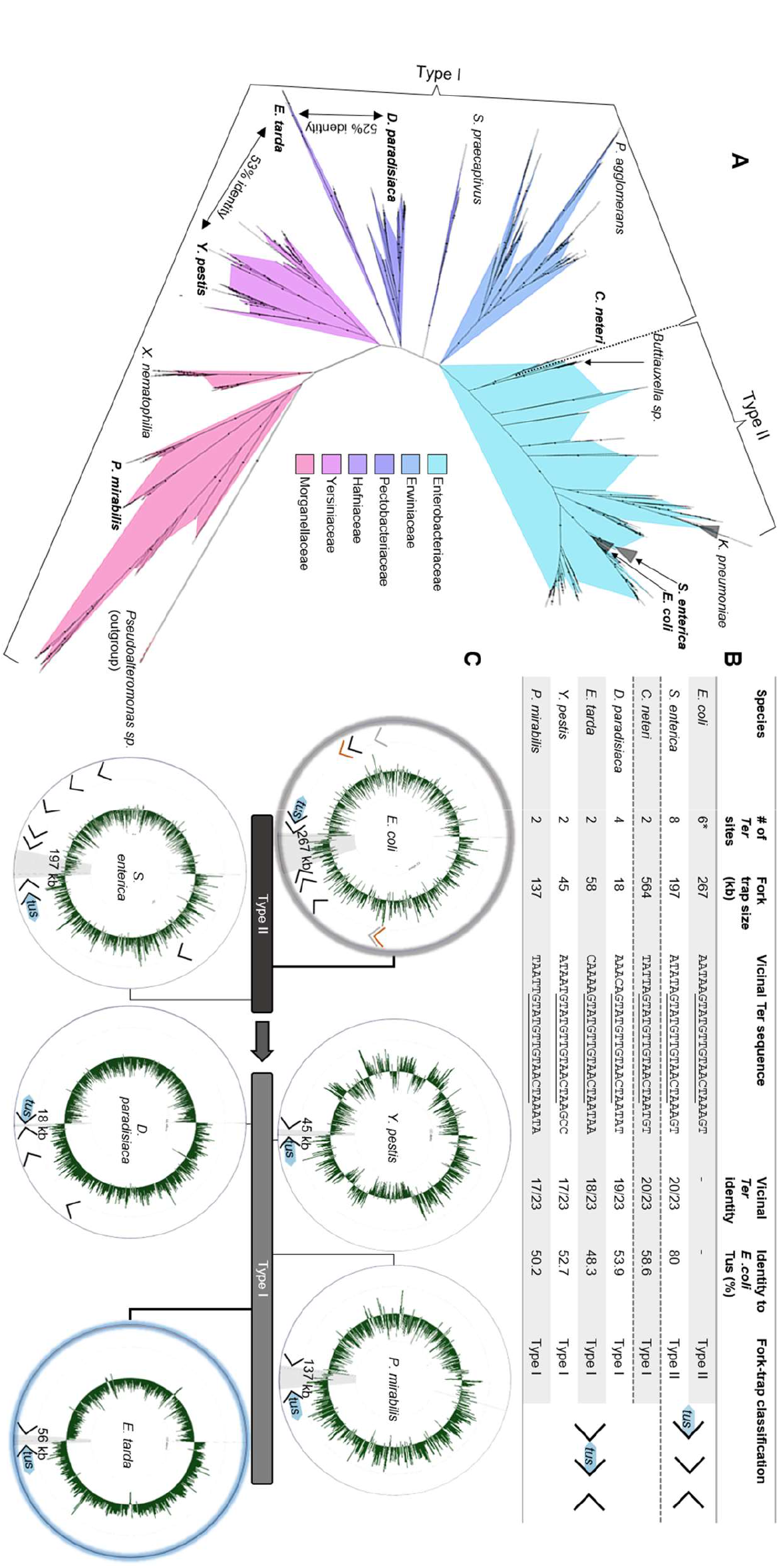
Phylogenetic analysis of Tus orthologs and fork trap architecture in Enterobacterales. (A) Unrooted phylogenetic relationship of ∼2500 Tus protein sequences using InterPro entries (IPR008865) highlighting the transition of a simple type I to complex type II fork trap architecture which occurs at *Cedecea*. (B) Chromosomal fork trap characteristics and classification for selected species (See SI Figure 7 & SI Table 1 for their graphical representations and the complete table of species). Fork trap size (kb) corresponds to the distance between the two innermost *Ter* sites of opposite polarity. Underlined bases represent a continuous identical sequence shared between all *Ter* sequences vicinal to *tus* starting at the GC(6) base-pair. (C) The different types of replication fork trap architecture in Enterobacterales.

In *Salmonella enterica*, a close relative of *E. coli*, the left chromosomal arm contains five *Ter* sites, while the right chromosomal arm contains only three *Ter* sites in opposite orientation (SI Figure 7A). The Tus protein and vicinal *Ter* sequence identities (i.e. corresponding to the *E. coli* Tus protein and *TerB*) were found to be 80% and 87% respectively. The distance between the innermost *Ter* sites (197 kb) is significantly reduced in the *Salmonella* replication fork trap. In more distantly-related bacteria, such as *Dickeya paradisiaca* and *Proteus mirabilis*, despite the high sequence identity of their respective *Ter* sequences vicinal to *tus* (83% for both), a reduction in the number of *Ter* sites as well as a narrowing of the fork trap (i.e. the distance between the innermost *Ter* sites), was commonly seen (Figure 3B). Most striking was that the innermost *Ter* site upstream of the *tus* gene (i.e. corresponding to the *E. coli TerC*) was no longer present in these species (Figure 3B). To our surprise, all replication fork traps that we characterised outside the Enterobacteriaceae family lacked the innermost *Ter* site corresponding to the *TerC* in *E. coli*. In these genomes (*D. paradisiaca*, *P. mirabilis, X. nematophilia*, *E. tarda* and *Yersinia pestis*), the innermost *Ter* site is the one vicinal to *tus* gene in that cluster (SI Figure 7). We thus propose a new replication fork trap classification based on their architecture where a type I replication fork trap has one of its innermost *Ter* sites vicinal to *tus* (Figure 3B). Accordingly, the *E. coli* and *S. enterica* genomes contain type II replication fork traps. All type I replication fork traps that we identified outside of the enterobacteriacae family are significantly narrower than the type II traps (Figure 3B). This is most evident in *D. paradisiaca* for which the innermost *Ter* inter-distance is just 18 kbp.

In *Edwardsiella tarda* this pattern of simplification culminated into a narrow and perfectly symmetrical replication fork trap diametrically opposite the *oriC*, consisting of two unique *Ter* sequences (Figure 3C). *E. tarda Ter1* and *Ter2* are only 56 kb apart and equidistantly located on either side of the hypothetical terminus site (Figure 4A). The next *Ter-*like sequence within this genome has only 65% identity to *Ter1* with a high level of degeneracy in the core sequence (see pseudo-*Ter3*, SI Figure 7F) and would oppose an o*riC*-initiated replication fork. Most importantly, the midpoint between *Ter1* and *Ter2* (∼1846 kb, *cf* Figure 4B) coincides almost perfectly with the sharp GC-skew flip (∼1847 kb) and *dif* (∼1843 kb) suggesting that they are being used equally as replication fork barriers. *E. tarda Ter1* shares 78% sequence identity to *E. coli TerB* (Figure 3B). In contrast, *E. coli TerC* shares only 74% sequence identity to *TerB* immediately suggesting that the mechanism of polar DNA replication fork arrest in *E. tarda* also involves formation of a locked Tus-*Ter* complex.

**Figure 4:**
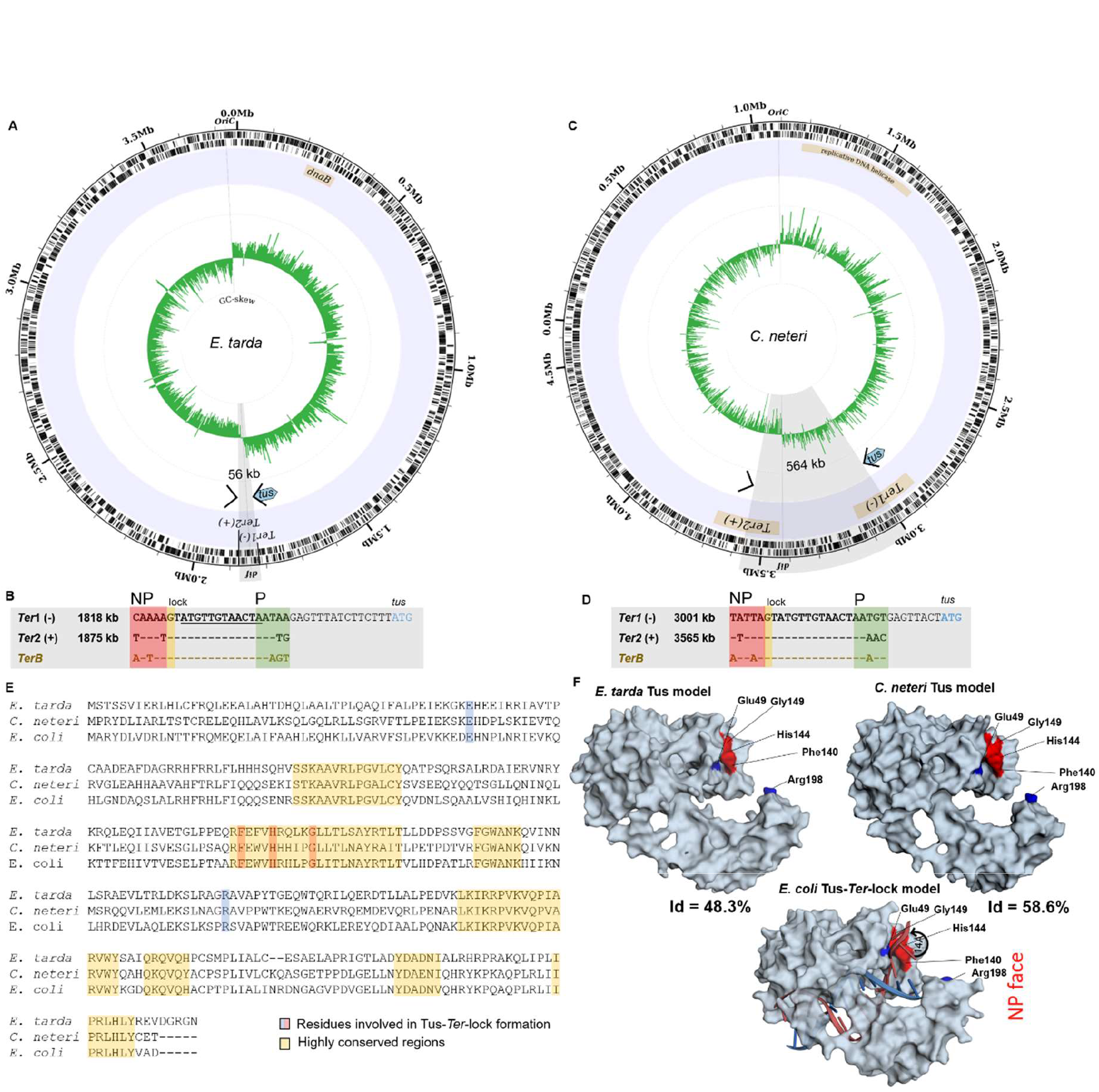
Prototypical type I replication fork trap. (A) Circular representation of *E. tarda* (strain EIB202) chromosome. Illustrated from the outside to the centre of the circle: forward and reverse genes, labelled genomic location of identified *Ter* sites involved in DNA replication termination, simplified annotation of the termination fork trap utilised, GC-skew over a 5000 bp moving window. The sharp GC-skew switches polarity at the replication origin and between the two identified *Ter* sites near *dif*. (B) Sequence alignment and genomic locations of the *E. tarda Ter* sites and *TerB* from *E. coli*. *Ter1* is located slightly upstream of the start site (ATG) of the *tus* gene similar to *TerB* in *E. coli.* The strictly conserved 12 bp core sequence is underlined and the G(6) base complementary to C(6) is highlighted in yellow. *NP: non-permissive face (red), P: permissive face (green).* (C) Circular representation of *C. neteri* (strain ND14a) chromosome. (D) Sequence alignment and genomic locations of the *C. neteri Ter* sites and *TerB* from *E. coli*. (E) Tus protein sequence alignment with highlighted conserved residues. (F) Comparison of the *E. coli* Tus-*Ter*-lock complex 3D structure (PDB 2I06) and the modelled structure of *E. tarda* and *C. neteri* Tus proteins using SWISS-MODEL. The essential amino acid residues in the cytosine binding pocket are indicated. The theoretical isoelectric points of *E. coli*, *E. tarda* and *C. neteri* Tus are 9.57, 9.67 and 9.31 respectively.

In our efforts to investigate the evolutionary divergence of the type I/II fork trap, we identified a unique group of *Cedecea* species within the Enterobacteriaceae family that uses a type I replication fork trap system with only two oppositely-oriented *Ter* sites for *C. neteri* (Figure 3B & Figure 4C-D). Against our expectations, the fork traps within this rare genus of bacteria were the widest (507-564 kb) of all investigated bacteria and the GC-skew switch (∼3420 kb) in *C. neteri* fitted unambiguously with the *dif* location (∼3424 kb). Of note, the *dif* site and GC-skew are located diametrically opposite the *oriC*, while the fork trap consisting of *Ter1* (3001 kb) and *Ter2* (3565 kb) is not.

While all vicinal *Ter* sequences that we examined are highly homologous (74-83% identity to *E. coli TerB)* and include the crucial C(6), and there is little doubt that the Tus orthologs from *S. enterica*, *Y. pestis* and *P. mirabilis* are able to arrest a replication fork at *TerB* (53); the competency of *C. neteri, D. paradisiaca*, *X. nematophilia* and *E. tarda* Tus orthologs (46-59% identity to *E. coli* Tus) to form a locked complex is not clear (Figure 3B). To examine if these Tus orthologs are competent in forming a locked complex (23), we verified that the residues that make a critical interaction with the C(6) base are strictly conserved (Figure 4E). It is apparent that *C. neteri* and especially *E. tarda* Tus, despite having one of the lowest identity score with the *E. coli* Tus sequence (48%), should be fully competent in forming locked complexes with their *Ter* sequences. Furthermore, model structures of *C. neteri* and *E. tarda* Tus showed no major differences in their respective cytosine binding pockets when compared to *E. coli* (Figure 4F) supporting formation of a highly efficient Tus-*Ter*-lock complex in both species. The sharp GC-skew flip midway between *Ter1* and *Ter2* suggests that replication forks rarely break-through the fork trap dyad in *E. tarda*. However, in *C. neteri*, it seems rather unlikely that *Ter1* and *Ter2* are being utilised to arrest replication forks.

## DISCUSSION

### A simplified type II replication fork trap in *E. coli*

Since the discovery of the first *Ter* sites and Tus coding sequence in *E. coli*, additional *Ter* sites were identified simultaneously expanding the size of the replication fork trap and increasing the perceived complexity of DNA replication termination. The systematic analyses of individual *Ter* sites both *in vitro* and *in vivo* with respect to their affinity and kinetics for Tus, ability of forming a TT-lock structure as well as their position and orientation within the bacterial genome have provided a wealth of information as to how this seemingly simple protein-DNA interaction impedes replication forks. In fact, Tus-*Ter* has become one of the best-understood protein-DNA complexes, leading to the development of a variety of biotechnological applications (54–57). Yet, we are only just starting to understand the modus operandi of Tus *in vivo*. Duggin and Bell showed evidence of a simple replication fork trap involving just *TerA, B and C* under normal bacterial concentrations of Tus (7). We found that the observed distribution of fork arrest events at these sites fits with the terminus GC-skew flip in the *E. coli* genome.

Taken together, our findings allow us to propose a simplified replication fork trap *in E. coli* consisting of just six *Ter* sites (three in each cluster) and support the notions that: (i) Tus binds preferentially to the high affinity *Ter* sites *in vivo*; (ii) Tus-bound *TerC* and *TerA* are sufficient to block replication forks progressing towards their non-permissive face; (iii) *TerB* is most likely only used when a replication fork passes through an unbound *TerC*; (iv) replication forks are unlikely to reach the outer *Ter* sites; (v) *TerH, I and J* are unlikely to be bound at natural Tus concentrations, are unable to block replication forks and thus cannot be considered as functional *Ter* sites.

While the roles of *TerA*, *B*, and *C* are now clear, the need for *TerD*, and particularly the distant *TerE* and *TerG* in the *E. coli* genome still remains somewhat enigmatic. If we consider that a single genomic insertion of *TerB* in the non-permissive orientation at SIR5.6 is not viable despite being the furthest from *tus* and the low natural abundance of Tus (58), this would support the notion that *TerD, E* and *G* although bound by Tus are rarely used to arrest replication forks.

### The prototypical type I DNA replication fork trap

The *E. coli* Tus-*Ter* mediated replication fork arrest mechanism has been intensely scrutinized in an attempt to better understand the final step in bacterial DNA replication. However, it now appears that the type II replication fork trap which is mostly found in Enterobacteriaceae is more of an exception or even an anomaly with respect to its many redundant *Ter* sites and their wide spread around the chromosome. It seems that the complexity of the type II fork trap in *E. coli* has merely distracted scientists from capturing the elegance and simplicity of the type I system in other Enterobacterales. Nevertheless, the work on the *E. coli* Tus-*Ter* complex was instrumental to decipher the unique TT-lock mechanism (10) which still stands true and is seemingly conserved in all *tus*-harboring bacteria.

Here, we show that the architecture and complexity of replication fork traps vary significantly across *tus*-harbouring bacteria. Yet, the two distinct classes of fork traps contain highly conserved *Ter* sequences despite moderate identity scores between Tus sequences. In the narrow type I fork trap, the *Ter1* vicinal to *tus* acts as a primary *Ter* site to arrest an incoming DnaB helicase travelling toward its non-permissive face (Figure 3B). The sharp terminal GC-skew switch observed in *E. tarda* strongly suggests that replication forks do not break through the *Ter* sites, and advocates against the need for back-up *Ter* sites. We propose that the narrow *E. tarda* fork trap consisting of only two *Ter* sites diametrically opposite of the *oriC* represents a prototypical type I replication fork trap.

In type II fork traps, the *Ter* vicinal to *tus* is rarely used as it is in second position from the terminus (*cf TerB* and *TerC* in *E. coli*, Figure 1A). We initially suspected that the large distance between *TerB* and *TerA* could have been the selective driver for acquiring an additional *Ter* site closer to the terminus and *dif* site in *E. coli* rather than the inherent need for a back-up system due to inefficient TT-lock formation that has previously been reported (14–16). However, our discovery of an unusually wide type I fork trap with a GC-skew switch fitting with the *dif* site in *Cedecea* species advocates against this possibility. Yet, it is unclear whether replication forks would even reach a blocking Tus-*Ter* complex in these bacteria as the GC-switch occurs diametrically opposite to the origin at the *dif* locus (Figure 4C).

### Conclusion and perspective

Our *in silico* data support the notion that all bacteria harbouring a type I fork trap use a Tus-*Ter* interaction that is competent in arresting an incoming replication fork by producing a TT-lock. We have discovered several Tus-dependent bacteria that do not require redundant *Ter* sites. The Tus-*Ter* interactions in these bacteria should have increased binding affinities and lock strengths due to the absence of back-up *Ter* sites in the fork trap. As such, it would be interesting to examine what the kinetic and thermodynamic parameters in these orthologous Tus-*Ter* systems are? Further examination of these prototypical fork trap systems will certainly help dissect the essential features and requirements of the unique Tus-*Ter* interaction.

Moving forward, a deeper understanding of the overall prevalence of the type I fork trap in bacteria is warranted to gain insight into the biological drivers that require or eliminate the need for redundant *Ter* sites in type II replication fork traps. Initially, our data might have suggested that in *E. coli* the distance of *TerB* from the terminus is not optimal for efficient replication termination and thus an additional *TerC* site with increased fork arrest activity was required to narrow the fork trap near the *dif* site. However, further data mining and the discovery of the extremely wide type I fork traps found in *Cedecae* species do not seem to support that hypothesis. Comparative examination of narrow versus wide type I fork traps could thus be key to shed further light on the evolutionary drivers for this system.

It is clear that the prototypical type I replication fork trap system will provide a great pedagogical tool for teaching DNA replication termination in curricula dealing with the central dogma in molecular biology. Furthermore, the development of a number of high-throughput proteomic and bioinformatic tools prompted by investigations into the *E. coli* Tus-*Ter* interaction, will no doubt facilitate further studies of orthologous systems. Indeed, Tus orthologs with different *Ter* binding-affinities would be very useful to develop finely tuneable assays to study DNA replication and transcription perturbation effects (54) and other biotechnologies (5, 56).

While the prototypical type I system clearly demonstrates that there is no inherent requirement for a back-up system to trap replication forks in the terminus region, the existence of both very narrow and very wide replication fork traps is puzzling. The wide type I fork traps found in *Cedecae* suggest that here, replication stalling activity may not be a primary purpose. As such, further comparative studies will be critical to fully decipher the mechanism of DNA replication termination and particularly the intersection between *dif* sites and fork traps as well as possible additional roles of Tus-*Ter e.g.* in chromosomal segregation (24). Finally, the diversity of type I and type II fork traps with respect to the number of *Ter* sites and their narrow or wide distribution begs the question as to what the evolutionary drivers for such variety are? Further examination of both wide and narrow type I replication fork traps will undoubtedly be instrumental to fully understand the replication fork trap and its possible interactions with other factors essential to DNA replication termination.

### DATA AVAILABILITY STATEMENT

ChIP-Seq and genome data have been deposited in GEO with the accession number GSE163680.

### FUNDING

The research was supported in part by the National Science Foundation Graduate Research Fellowship under Grant No DGE-1110007, the National Security Science and Engineering Faculty Fellowship (FA9550-10-1-0169), and the Welch Foundation (F-1654).

## Supporting information

Additional Resources

