## Supplementary Information for "Defining the prototypical DNA replication fork trap in bacteria"

### MATERIALS AND OTHER RESOURCES

| REAGENT or RESOURCE | SOURCE | IDENTIFIER |
|---|---|---|
| Antibodies | | |
| Chicken anti-GFP IgY | Abcam | ab92456 |
| HRP-conjugated goat anti-IgY (Jackson 103-035-155) | Jackson ImmunoResearch Laboratories | 103-035-155 |
| Bacterial Strains | | |
| *E. coli* KRX | Promega | Cat#: L3002 |
| BL21(*DE3*)RIPL | Stratagene | Cat#: 230280 |
| *Dickeya paradisiaca* (strain Ech703) | RefSeq | NC_012880 |
| *Edwardsiella tarda* (strain EIB202) | RefSeq | NC_013508 |
| *Proteus mirabilis (strain HN2p)* | RefSeq | NZ_CP046048 |
| *Xenorhabdus nematophila* (strain ATCC 19061) | RefSeq | NC_014228 |
| *Salmonella typhimurium* (strain LT2) | RefSeq | NC_003197 |
| *Escherichia coli* (strain K12 substr. MG1655) | RefSeq | U00096 |
| *Cedecea neteri* (strain ND14a) | RefSeq | NZ_CP009459 |
| Recombinant DNA | | |
| Plasmid: pPMS1259 | Schaeffer Lab | (1) |
| Chemicals, Peptides, and Recombinant Proteins | | |

| | | |
|---|---|---|
| $His_6$-Tus-GFP | Schaeffer Lab | N/A |
| SIGMAFAST™ 3,3′ -Diaminobenzidine tablets | Sigma | d4418 |
| SensiMix SYBR & fluorescein mastermix | Bioline | QT615-05 |
| Critical Commercial Assays | | |
| NEBNext Ultra DNA library preparation kit | New England BioLabs | E7370S |
| QuantiFluor® dsDNA System | Promega | E2670 |
| Rapid Sequencing protocol (FLO-MIN106 R9 MinION) | Oxford Nanopore | SQK-RAD004 |
| Deposited Data | | |
| ChIP-Seq data set and KRX assembly | NCBI GEO | Accession: GSE163680 |
| Oligonucleotides | | |
| See Supplementary Data for full list of sequences and genomic loci for amplification of *oriC* and *Ter* regions by qPCR | | |
| Software and Algorithms | | |
| MinKNOW | Oxford Nanopore Technologies | https://github.com/nanoporetech/minknow_api |
| Trimmomatic | (2) | http://www.usadellab.org/cms/?page=trimmomatic |

| | | |
|---|---|---|
| Porechop | (3) | https://github.com/rrwick/Porechop |
| Flye | (4) | https://github.com/fenderglass/Flye |
| Racon | (5) | https://github.com/isovic/racon |
| Pilon | (6) | https://github.com/broadinstitute/pilon |
| Quast | (7) | http://quast.sourceforge.net/quast |
| Prokka | (8) | https://github.com/tseemann/prokka |
| Bowtie2 | (9) | http://bowtie-bio.sourceforge.net/bowtie2/index.shtml |
| Samtools | (10) | http://www.htslib.org/ |
| Circleator | (11) | http://jonathancrabtree.github.io/Circleator/ |
| genomeCoverageBed | (12) | https://bedtools.readthedocs.io/en/latest/content/tools/genomecov.html |

| | | |
|---|---|---|
| blastn | (13) | https://ftp.ncbi.nlm.nih.gov/blast/executables/blast+/LATEST/ |
| Interactive Genomics Viewer (IGV) | (14) | http://software.broadinstitute.org/software/igv/ |
| EzMol Molecular display wizard | (15) | http://www.sbg.bio.ic.ac.uk/ezmol/ |
| InterPro Protein Data Bank | (16) | https://www.ebi.ac.uk/interpro/ |
| iTol | (17) | https://itol.embl.de/ |
| RAxML | (18) | https://cme.h-its.org/exelixis/web/software/raxml/ |
| MUSCLE | (19) | http://www.drive5.com/muscle/downloads.htm |
| ImageJ | (20) | https://imagej.nih.gov/ij/index.html |
| GraphPad 8 | GraphPad Software | https://www.graphpad.com/scientific-software/prism/ |

### CONTACT FOR REAGENT AND RESOURCE SHARING

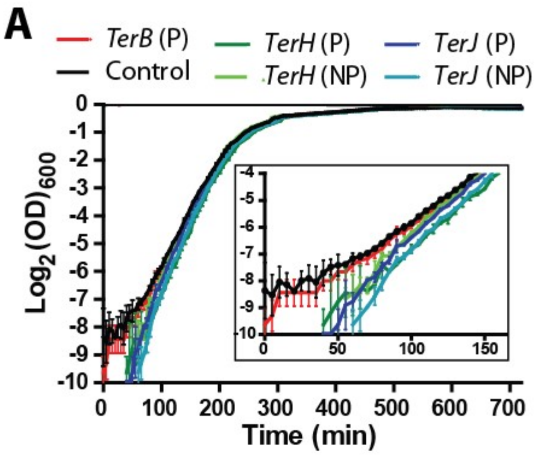

**SI Figure 1: Effect of ectopic Ter sites on the growth rate of E. coli BL21(DE3).** *TerB*, *TerH* and *TerJ* were inserted ~ 930 kpb downstream of *oriC* in the permissive (P) or non-permissive (NP) orientation. (A) Growth rates were measured in independent triplicates. Error bars represent SD. A culture of wild type BL21(DE3) was grown as a control. Growth rates were determined from the slopes of the linear regressions performed between 100 and 210 minutes (see Table 1 in the main text). Doubling time ($T_D$) was calculated as 1/growth rate (n=3, except for TerH (NP), n=2). Reproduced with permission from Moreau, PhD thesis, James Cook University (2013). Thesis can be downloaded using the following link: ***https://researchonline.jcu.edu.au/31903/1/31903_Moreau_2013_thesis.pdf***).

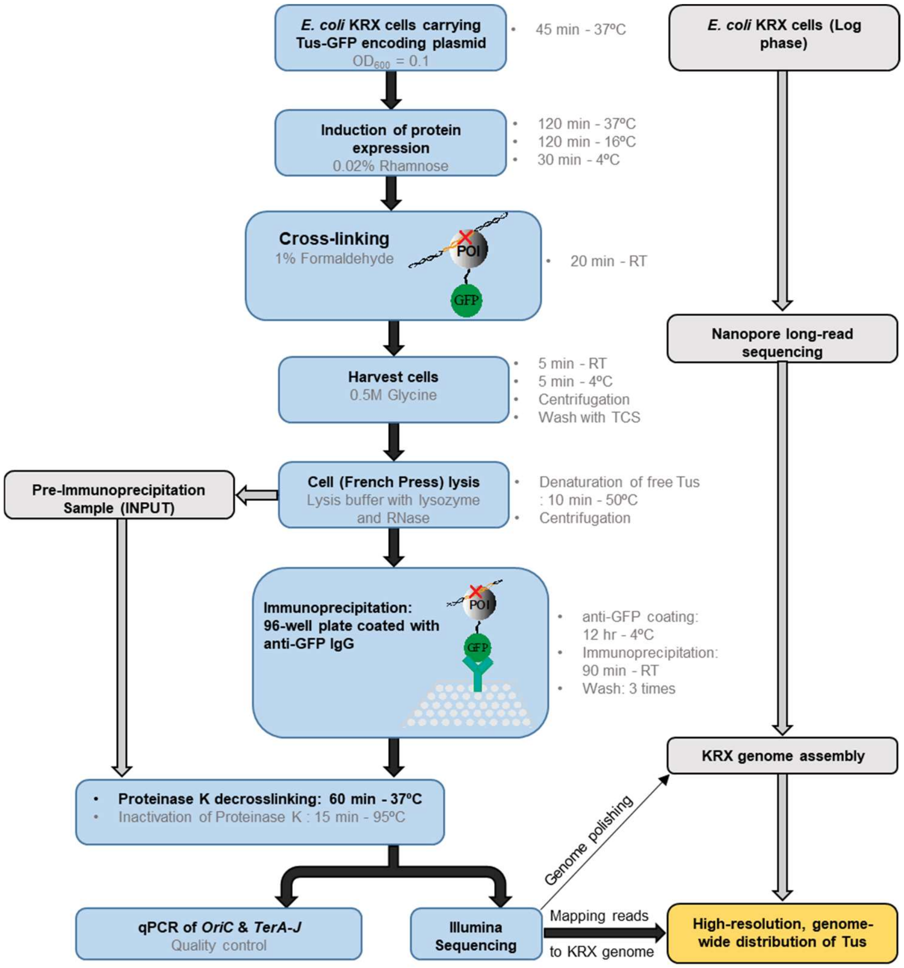

**SI Figure 2: ChIP-qPCR and ChIP-Seq process using a 96-well plate format coated with anti-GFP IgG, and genome assembly for *E. coli* KRX strain.** See Star Methods section for detailed procedures.

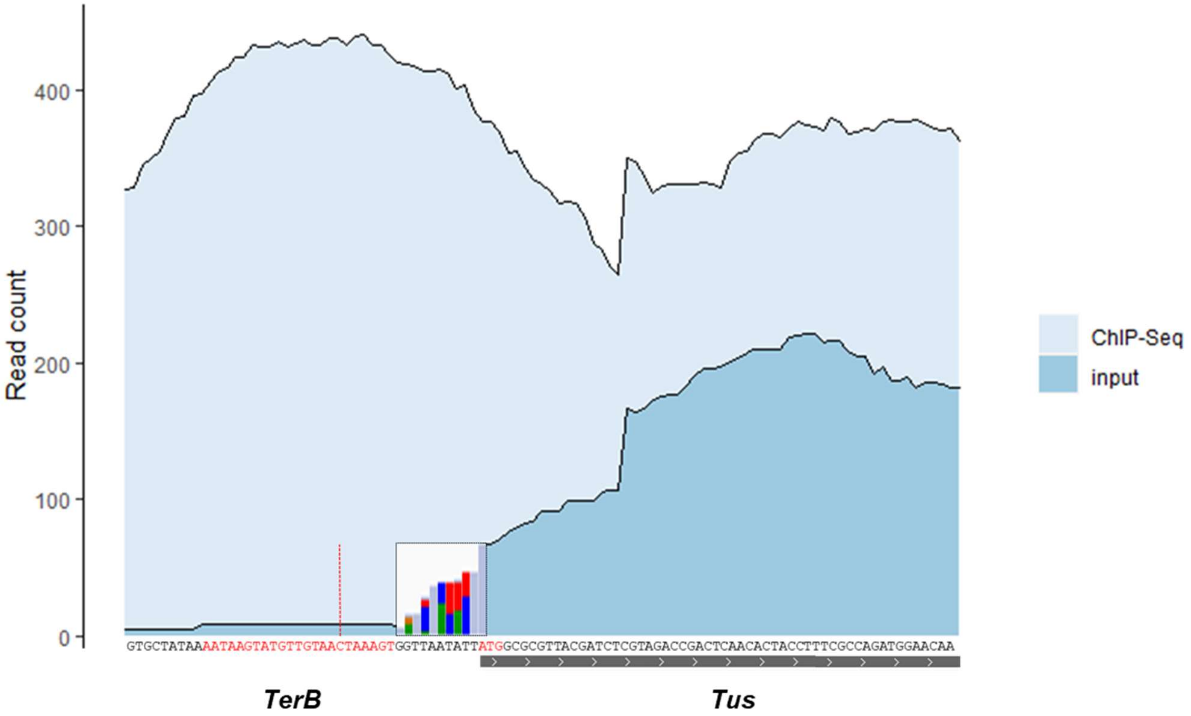

**SI Figure 3: Nucleotide read count at genomic *TerB* and *tus* gene loci for immunoprecipitated DNA (ChIP-Seq) and non-immunoprecititated DNA (Input).** The boxed bar chart between *TerB* and *tus* shows an ambiguous 10 nucleotide sequence with partial identity between the plasmid and genome sequences upstream the start codon (ATG) of the *tus* sequence. The data show that the high read count originating from the plasmid *tus* sequence (i.e. misaligned to the genomic *tus* locus) does not bias the read count at *TerB*. Based on the average *tus* read count the plasmid number is ~100-200/bacteria. The dashed red line indicates the C(6) position in *TerB* critical for the formation of the Tus-*Ter*-lock structure.

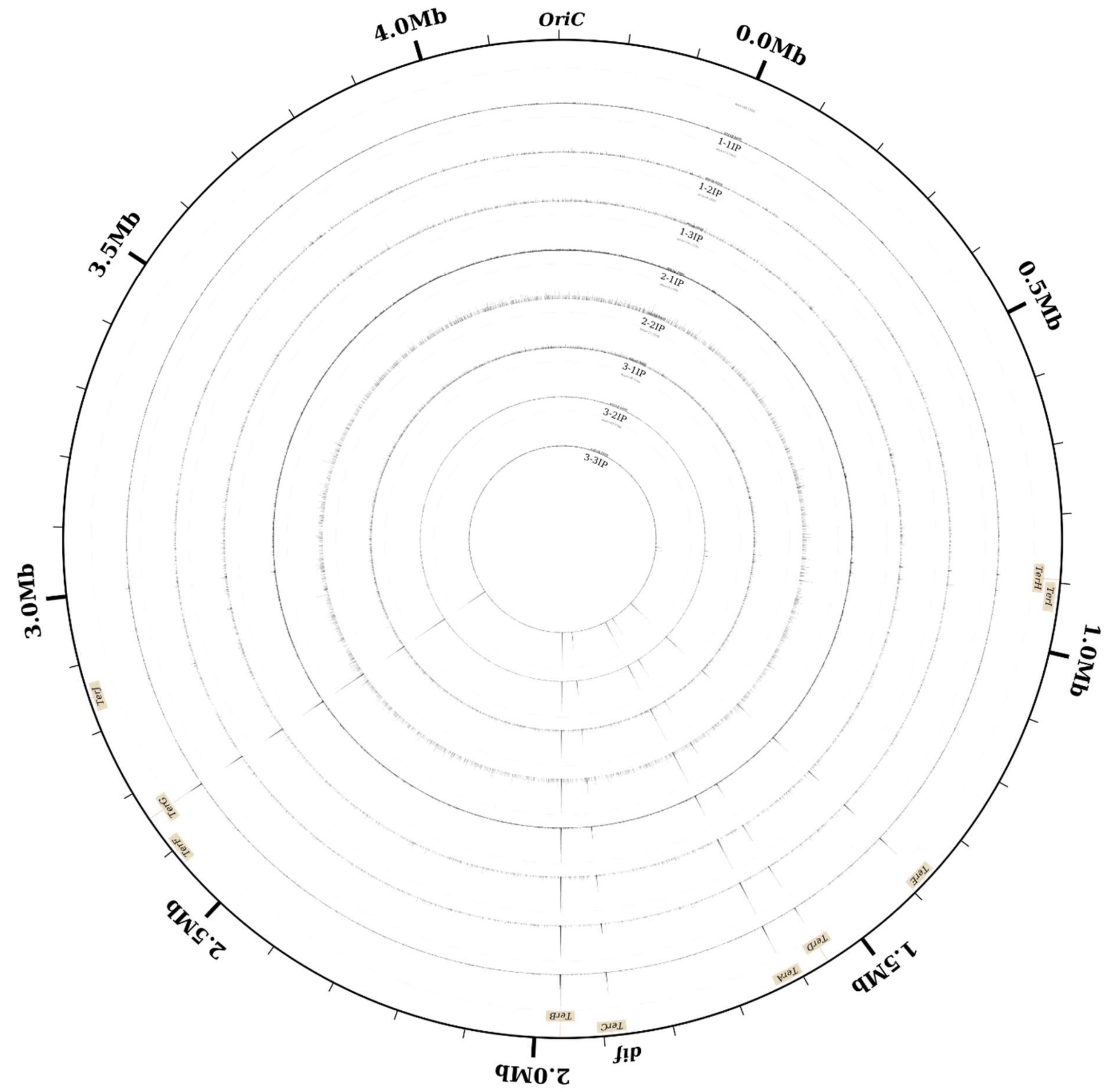

**SI Figure 4: Individually mapped ChIP-Seq coverage for experimental and biological replicates (n = 8).** Three biological replicates of immunoprecipitated DNA (ChIP) are shown each consisting of 2-3 technical replicates as indicated. The difference in peak height between different *Ter* sites is consistent despite varying depths of sequencing between replicates. See Figure 2 for pooled data.

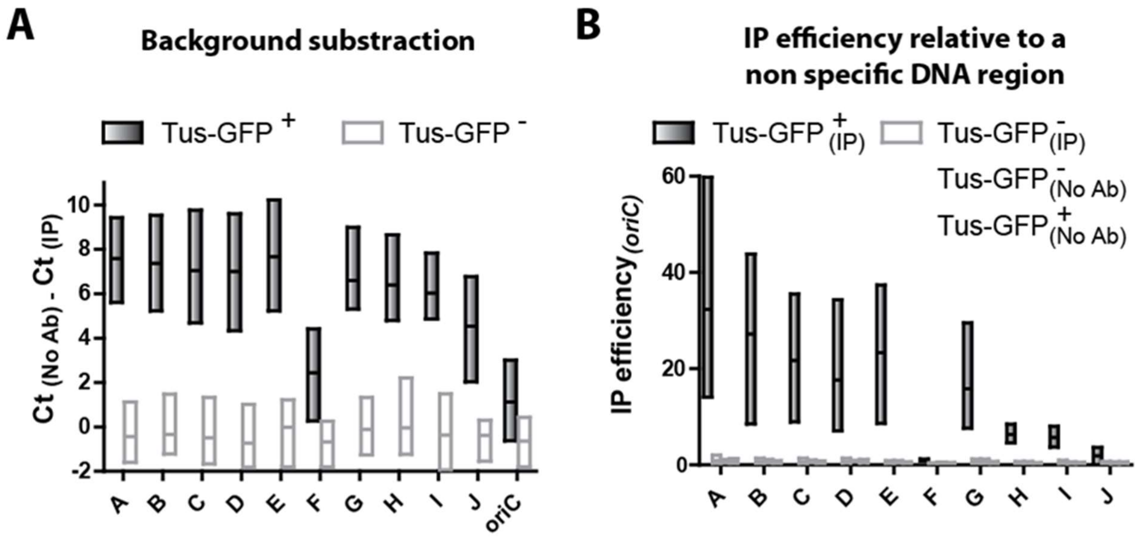

**SI Figure 5: Distribution of Tus-GFP on *Ter* sites in *E. coli* KRX cells by ChIP-qPCR**. (A) Difference in Ct-values between immunoprecipitated DNA (IP) and background control experiments in absence of anti-GFP (No Ab) obtained for Tus-GFP$^+$ and Tus-GFP$^-$ control KRX cells. (B) IP efficiency of *Ter* sites relative to a non-specific *oriC* region obtained for Tus-GFP$^+$ and Tus-GFP$^-$ control KRXcells in the presence (IP) or absence of anti-GFP IgG antibody (No Ab). Floating bars represent minimum, maximum and mean values. Reproduced with permission from Moreau, PhD thesis, James Cook University (2013). Thesis can be downloaded at: https://researchonline.jcu.edu.au/31903/1/31903_Moreau_2013_thesis.pdf.

A

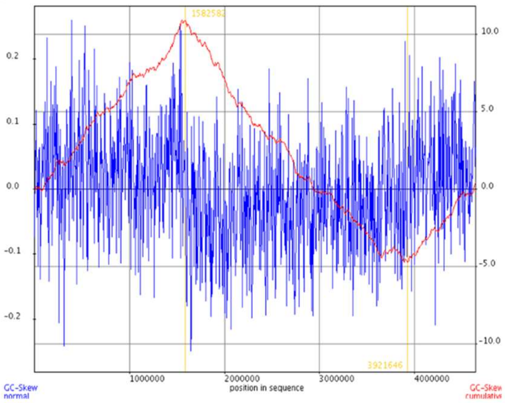

1,582,582

B

| | *TerA* | *TerB* | *TerC* | *dif* | GC skew transition midpoint |
|---|---|---|---|---|---|
| Location in *E. coli* genome (bp) | 1,341,745 | 1,684,226 | 1,609,157 | 1,590,777 | 1,582,582 |
| *In vivo* replication arrest (%) | 0.19 | 0.14 | 0.85 | - | - |
| Fraction of arrest Hypothesis 1 | 0.161 | 0.119 | 0.72 | - | - |
| Fraction of arrest Hypothesis 2 | 0.183 | - | 0.817 | - | - |
| Fraction of arrest Hypothesis 3 | - | - | - | 1 | - |
| | | | | | **Distance from GC skew (bp)** |
| **Predicted GC-inflection Hypothesis 1 (bp)** | 1,575,037 | | | | -7545 |
| **Predicted GC-inflection Hypothesis 2 (bp)** | 1,560,221 | | | | -22361 |
| **Predicted GC-inflection Hypothesis 3** | 1,590,777 | | | | 8195 |

**SI Figure 6**: (A) GC skew transition midpoint calculated with a 1000 bp sliding window for *E. coli* MG1655 used in the *in vivo* replication arrest study by Duggin and Bell, (2009). (B) Hypothetical GC skew transition midpoint loci using various fork arrest scenarios based on the ensemble and fractional distribution of replication fork collision loci at functional *Ter* sites. Only *Ter* sites with significant replication fork arrest activity (*TerA*, *TerB* and *TerC*) are included. *Dif* site is also shown for comparison. Locations of *TerA-C* and *dif* in *E. coli* MG1655 are indicated in the table.

**Hypothesis 1:** Replication fork arrest occurs with the fractional distribution of Y forks reported by Duggin and Bell, 2009.

**Hypothesis 2:** Replication fork arrest occurs at *TerA* and *TerC* with equal fractional distribution i.e. fork arrest only occurs at *TerA* and *TerC* and are never breached.

**Hypothesis 3:** Replication fork arrest occurs at *dif* site

**Conclusion:** the terminal GC-skew swith derived from hypothesis 1 (i.e. 1,575,037 bp) involving *TerA-C* deviates the least from the switch point derived from a sliding 1,000 bp cumulative GC-skew (i.e. 1,582,582 bp) by only 7.5 kbp. It is important to note that the *dif* site is located 8 kbp from the terminal GC-skew switch point on the other chromosomal arm.

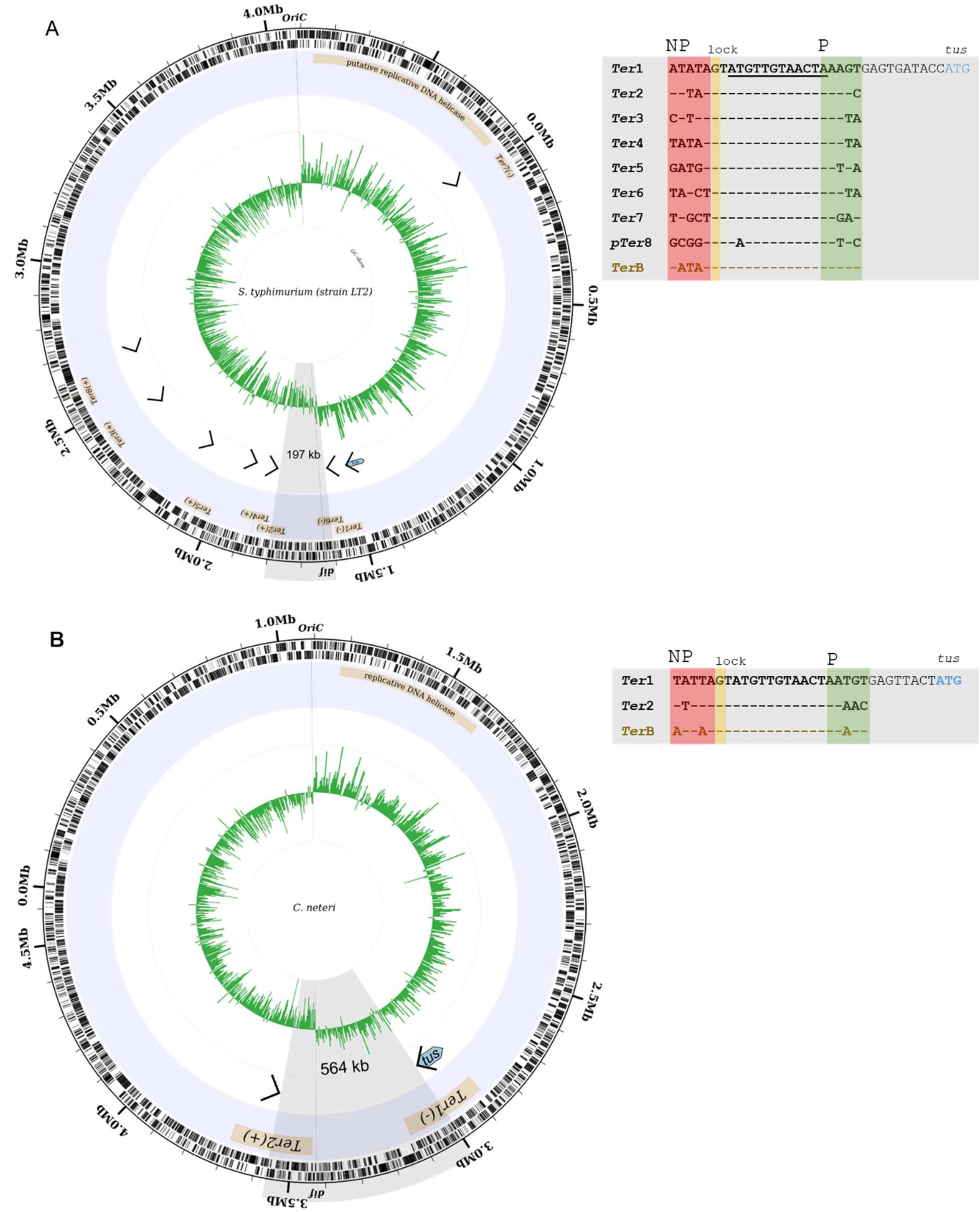

A
4.0Mb
OriC
3.5Mb
0.0Mb
3.0Mb
0.5Mb
2.5Mb
1.0Mb
2.0Mb
1.5Mb
dif
putative replicative DNA helicase
S. typhimurium (strain LT2)
197 kb
NP lock P tus
Ter1 ATATAGTATGTTGTAACTAAAGTGAGTGATACCATG
Ter2 --TA---------------C
Ter3 C-T----------------TA
Ter4 TATA---------------TA
Ter5 GATG---------------T-A
Ter6 TA-CT--------------TA
Ter7 T-GCT--------------GA-
pTer8 GCGG---A-----------T-C
TerB -ATA-------------------
B
1.0Mb
OriC
1.5Mb
0.5Mb
2.0Mb
0.0Mb
4.5Mb
2.5Mb
4.0Mb
3.0Mb
3.5Mb
dif
replicative DNA helicase
C. neteri
564 kb
tus
Ter2(+)
Ter1(-)
NP lock P tus
Ter1 TATTAGTATGTTGTAACTAATGTGAGTTACTATG
Ter2 -T-----------------AAC
TerB A--A---------------A--

C

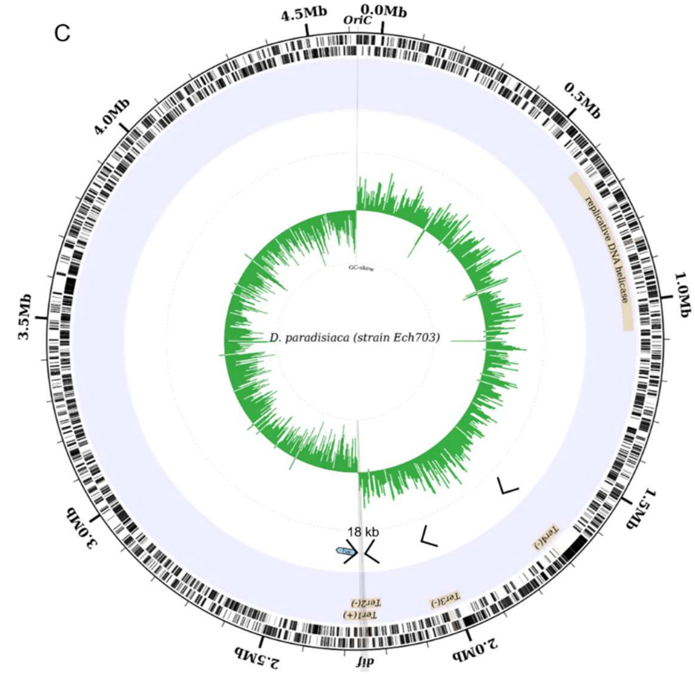

| | NP | lock | | P | tus |
|---|---|---|---|---|---|
| ***Ter1*** | AAACA | G | TATGTTGTAACT | AATAT | GGTGTTTCTATG |
| ***Ter2*** | C-TT- | - | ------------ | ---T- |  |
| ***Ter3*** | T--A- | - | ------------ | -T-TA |  |
| ***Ter4*** | GGC-- | - | A----------- | CCCG- |  |
| ***TerB*** | --TA- | - | ------------ | --AG- |  |

D

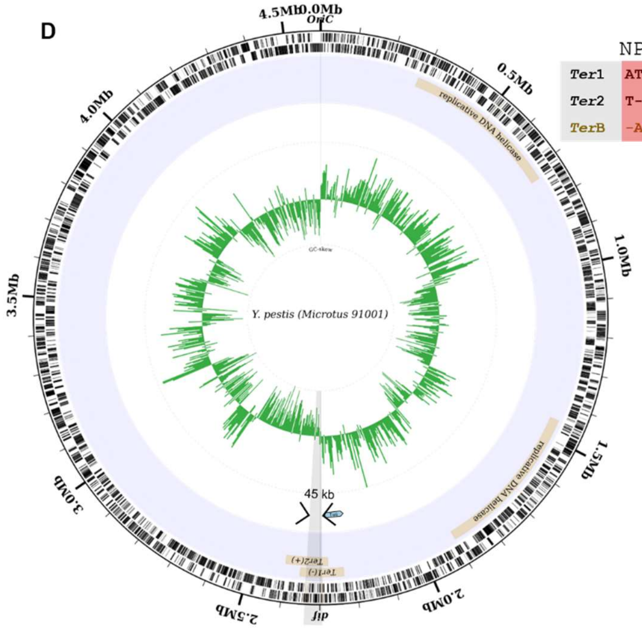

| | NP | lock | | P | tus |
|---|---|---|---|---|---|
| ***Ter1*** | ATAAT | G | TATGTTGTAACT | AAGCC | GCCTTATTTAGAAAAGAATATGAACAAGTATG |
| ***Ter2*** | T--C- | - | ------------ | --AAT |  |
| ***TerB*** | -AT-A | - | ------------ | --AGT |  |

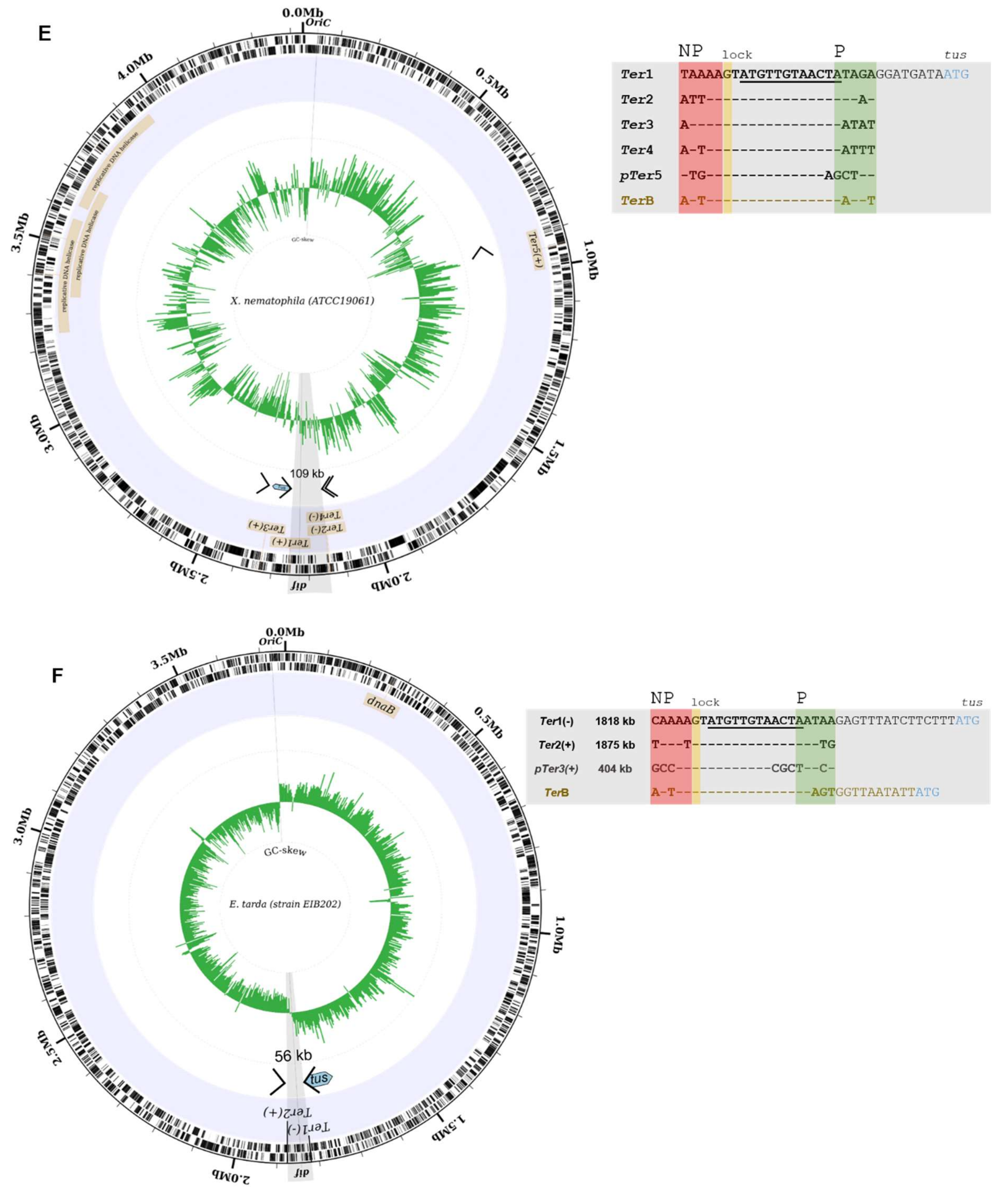

**SI Figure 7: Circular representation of (A) *Salmonella typhimurium, (B) Cedecae netari, (C) Dickeya paradisiaca, (D) Yersinia pestis (E) Xenorhabdus nematophila* and *(F) Edwardsiella tarda* chromosomes and their *Ter* sequences.** From the outside of the circle: Forward and reverse genes; genomic locations of identified *Ter* sites involved in DNA replication termination; and GC-skew with a 5000 bp sliding window. In each alignment, *Ter1* represents the *Ter* sequence adjacent to *tus* in the indicated chromosome followed by the RBS and start codon of *tus*. The G(6) complementary to C(6) is highlighted in yellow and the strictly conserved 12 bp core sequence is underlined. The non-permissive face (NP) is highlighted in red and the permissive face (P) is highlighted in green.

**SI Table 1: Chromosomal fork trap architecture and classification for selected bacteria.**

| Family | Species | # of *Ter* sites | Fork trap size (kb) | Vicinal *Ter* sequence | Vicinal *Ter* identity | Identity to *E .coli* Tus (%) | Fork-trap classification |
|---|---|---|---|---|---|---|---|
| Enterobacteriacae | *Escherichia coli* | 6* | 267 | AATAAGTATGTTGTAACTAAAGT | - | - | Type II |
| | *Salmonella typhimurium* | 8 | 197 | ATATAGTATGTTGTAACTAAAGT | 20/23 | 80 | Type II |
| | *Cronobacter dublinensis* | 9 | 47 | ATAAAGTATGTTGTAACTAATGT | 20/23 | 65 | Type II |
| | *Atlantibacter hermannii* | 8 | 43 | AAATAGTATGTTGTAACTAAAGG | 20/23 | 61.8 | Type II 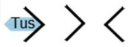 |
| | *Shimwellia blattae* | 5 | 72 | AATAAGCATGTTGTAACTAAAGA | 21/23 | 60 | Type II |
| | *Buttiauxella agrestis* | 5 | 297 | CTTTAGTATGTTGTAACTAATGG | 18/23 | 60.5 | Type II |
| | *Cedecea netari str. FDAARGOS* | 3 | 507 | CATTAGTATGTTGTAACTAAAGT | 21/23 | 59.2 | Type I |
| | *Cedecea neteri str. ND14a* | 2 | 564 | TATTAGTATGTTGTAACTAATGT | 20/23 | 58.6 | Type I |
| Erwiniaceae | *Pantoea agglomerans* | 4 | 246 | TTATAGTATGTTGTAACTATAAA | 16/23 | 55.4 | Type I |
| Pectobacteriaceae | *Sodalis praecaptivus* | 2 | 111 | GTATAGTATGTTGTAACTAATAG | 16/23 | 50.2 | Type I |
| | *Dickeya Paradisiaca* | 4 | 18 | AAACAGTATGTTGTAACTAATAT | 19/23 | 53.9 | Type I 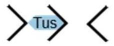 |
| Hafniaceae | *Edwardsiella tarda* | 2 | 58 | CAAAAGTATGTTGTAACTAATAA | 18/23 | 48.3 | Type I |
| Yersiniaceae | *Yersinia pestis* | 2 | 45 | ATAATGTATGTTGTAACTAAGCC | 17/23 | 52.7 | Type I |
| Morganellaceae | *Xenorhabdus nematophila* | 4 | 109 | TAAAAGTATGTTGTAACTATAGA | 19/23 | 46.3 | Type I |
| | *Proteus mirabilis* | 2 | 137 | TAATTGTATGTTGTAACTAAATA | 17/23 | 50.8 | Type I |

*Fork trap size corresponds to the distance between the two innermost Ter sites of opposite polarity expressed in kb. Underlined bases represent a continuous identical sequence shared between all Ter sequences vicinal to tus starting at the G(6).*
