## Additional Resources for "Defining the prototypical DNA replication fork trap in bacteria"

```
BLASTN 2.10.1+: Matrix: blastn matrix 1 -3,
Gap Penalties: Existence: 5, Extension: 2

Database: final_high_quality_krx_assembly.fasta
           1 sequences; 4,491,350 total letters
```

**Query= TerAF**

```
Length=24

>contig_2_pilon
Length=4491350

 Score = 48.1 bits (24),  Expect = 2e-07
 Identities = 24/24 (100%), Gaps = 0/24 (0%)
 Strand=Plus/Plus

Query  1        CAACCATTAAACCGATTCGCGGTC  24
                ||||||||||||||||||||||||
Sbjct  1619469  CAACCATTAAACCGATTCGCGGTC  1619492
```

**Query= TerAR**

```
Length=20

>contig_2_pilon
Length=4491350

 Score = 40.1 bits (20),  Expect = 3e-05
 Identities = 20/20 (100%), Gaps = 0/20 (0%)
 Strand=Plus/Minus

Query  1        AGTTGCGATTTCTCCCCTGG  20
                ||||||||||||||||||||
Sbjct  1619613  AGTTGCGATTTCTCCCCTGG  1619594
```

**Query= TerBF**

```
Length=22

>contig_2_pilon
Length=4491350

 Score = 44.1 bits (22),  Expect = 2e-06
 Identities = 22/22 (100%), Gaps = 0/22 (0%)
 Strand=Plus/Plus


Query  1        TTACCTCTGCCTGACACTACGC  22
                ||||||||||||||||||||||
Sbjct  1962934  TTACCTCTGCCTGACACTACGC  1962955
```

**Query= TerBR**

```
Length=23

>contig_2_pilon
Length=4491350
 Score = 46.1 bits (23),  Expect = 6e-07
 Identities = 23/23 (100%), Gaps = 0/23 (0%)
 Strand=Plus/Minus

Query  1        TGTTGAGTCGGTCTACGAGATCG  23
                |||||||||||||||||||||||
Sbjct  1963056  TGTTGAGTCGGTCTACGAGATCG  1963034
```

**Query= TerCF**

```
>contig_2_pilon
Length=4491350

 Score = 46.1 bits (23),  Expect = 6e-07
 Identities = 23/23 (100%), Gaps = 0/23 (0%)
 Strand=Plus/Plus

Query  1        CTGCATGTGGCACCTGTTAATGA  23
                |||||||||||||||||||||||
Sbjct  1887884  CTGCATGTGGCACCTGTTAATGA  1887906
```

**Query= TerCR**

```
Length=20

>contig_2_pilon
Length=4491350

 Score = 40.1 bits (20),  Expect = 3e-05
 Identities = 20/20 (100%), Gaps = 0/20 (0%)
 Strand=Plus/Minus

Query  1        GCTGTACGTCCGTTGTGCTA  20
                ||||||||||||||||||||
Sbjct  1888006  GCTGTACGTCCGTTGTGCTA  1887987
```

**Query= TerDF**

```
Length=25

>contig_2_pilon
Length=4491350

 Score = 50.1 bits (25),  Expect = 5e-08
 Identities = 25/25 (100%), Gaps = 0/25 (0%)
 Strand=Plus/Plus

Query  1        GGCATGATGTCGCGCttttttttATG  25
                |||||||||||||||||||||||||
Sbjct  1558486  GGCATGATGTCGCGCTTTTTTTATG  1558510
```

**Query= TerDR**

```
Length=25

>contig_2_pilon
Length=4491350

 Score = 50.1 bits (25),  Expect = 5e-08
 Identities = 25/25 (100%), Gaps = 0/25 (0%)
 Strand=Plus/Minus

Query  1        GGGTATTAAGGAGTATTCCCCATGG  25
                |||||||||||||||||||||||||
Sbjct  1558610  GGGTATTAAGGAGTATTCCCCATGG  1558586
```

**Query= TerEF**

```
Length=20

>contig_2_pilon
```

```
Length=4491350

 Score = 40.1 bits (20),  Expect = 3e-05
 Identities = 20/20 (100%), Gaps = 0/20 (0%)
 Strand=Plus/Plus

Query  1        GAAGTCGCCGTCTGGTTTAT  20
                ||||||||||||||||||||
Sbjct  1377410  GAAGTCGCCGTCTGGTTTAT  1377429
```

**Query= TerER**

```
Length=20

>contig_2_pilon
Length=4491350

 Score = 40.1 bits (20),  Expect = 3e-05
 Identities = 20/20 (100%), Gaps = 0/20 (0%)
 Strand=Plus/Minus

Query  1        TACGGCGGAAGTTAATGGTC  20
                ||||||||||||||||||||
Sbjct  1377581  TACGGCGGAAGTTAATGGTC  1377562
```

**Query= TerFF**

```
Length=21

>contig_2_pilon
Length=4491350

 Score = 42.1 bits (21),  Expect = 8e-06
 Identities = 21/21 (100%), Gaps = 0/21 (0%)
 Strand=Plus/Plus

Query  1        CACATCTTCGGGAGTCGGTTC  21
                |||||||||||||||||||||
Sbjct  2596624  CACATCTTCGGGAGTCGGTTC  2596644
```

**Query= TerFR**

```
Length=22

>contig_2_pilon
Length=4491350

 Score = 44.1 bits (22),  Expect = 2e-06
 Identities = 22/22 (100%), Gaps = 0/22 (0%)
 Strand=Plus/Minus

Query  1        GGTTGAGTGGTAAACGCTGCTG  22
                ||||||||||||||||||||||
Sbjct  2596754  GGTTGAGTGGTAAACGCTGCTG  2596733
```

**Query= TerGF**

```
Length=20

>contig_2_pilon
Length=4491350

 Score = 40.1 bits (20),  Expect = 3e-05
 Identities = 20/20 (100%), Gaps = 0/20 (0%)
```

```
 Strand=Plus/Plus

Query  1        CCAAGCGAGTACCCCACCAG  20
                ||||||||||||||||||||
Sbjct  2656294  CCAAGCGAGTACCCCACCAG  2656313
```

**Query= TerGR**

```
Length=23

>contig_2_pilon
Length=4491350

 Score = 46.1 bits (23),  Expect = 6e-07
 Identities = 23/23 (100%), Gaps = 0/23 (0%)
 Strand=Plus/Minus

Query  1        CACGGTTGTATGTTGATCTCCCA  23
                |||||||||||||||||||||||
Sbjct  2656435  CACGGTTGTATGTTGATCTCCCA  2656413
```

**Query= TerHF**

```
Length=24

>contig_2_pilon
Length=4491350

 Score = 48.1 bits (24),  Expect = 2e-07
 Identities = 24/24 (100%), Gaps = 0/24 (0%)
 Strand=Plus/Plus

Query  1       TGAAGGACAAACTGGAAACGCTGA  24
               ||||||||||||||||||||||||
Sbjct  895054  TGAAGGACAAACTGGAAACGCTGA  895077
```

**Query= TerHR**

```
Length=20

>contig_2_pilon
Length=4491350

 Score = 40.1 bits (20),  Expect = 3e-05
 Identities = 20/20 (100%), Gaps = 0/20 (0%)
 Strand=Plus/Minus

Query  1       CAGACTACCGCCACCACAAT  20
               ||||||||||||||||||||
Sbjct  895201  CAGACTACCGCCACCACAAT  895182
```

**Query= TerIF**

```
Length=22

>contig_2_pilon
Length=4491350

 Score = 44.1 bits (22),  Expect = 2e-06
 Identities = 22/22 (100%), Gaps = 0/22 (0%)
 Strand=Plus/Plus

Query  1       ATTGCTGGAACGGTTGATTGCG  22
               ||||||||||||||||||||||
Sbjct  920804  ATTGCTGGAACGGTTGATTGCG  920825
```

**Query= TerIR**

```
Length=20
>contig_2_pilon
Length=4491350

 Score = 40.1 bits (20),  Expect = 3e-05
 Identities = 20/20 (100%), Gaps = 0/20 (0%)
 Strand=Plus/Minus

Query  1       CTCGCCGTCTTTACGTAGCA  20
               ||||||||||||||||||||
Sbjct  920921  CTCGCCGTCTTTACGTAGCA  920902
```

**Query= TerJF**

```
Length=20

>contig_2_pilon
Length=4491350

 Score = 40.1 bits (20),  Expect = 3e-05
 Identities = 20/20 (100%), Gaps = 0/20 (0%)
 Strand=Plus/Plus

Query  1        GACGATACGACGCACCGATG  20
                ||||||||||||||||||||
Sbjct  2849492  GACGATACGACGCACCGATG  2849511
```

**Query= TerJR**

```
Length=22
>contig_2_pilon
Length=4491350

 Score = 44.1 bits (22),  Expect = 2e-06
 Identities = 22/22 (100%), Gaps = 0/22 (0%)
 Strand=Plus/Minus

Query  1        CTGGTGATGCCGAACATGGAAG  22
                ||||||||||||||||||||||
Sbjct  2849641  CTGGTGATGCCGAACATGGAAG  2849620
```

**Query= OriCF**

```
Length=22
>contig_2_pilon
Length=4491350

 Score = 44.1 bits (22),  Expect = 2e-06
 Identities = 22/22 (100%), Gaps = 0/22 (0%)
 Strand=Plus/Plus

Query  1        CGCACTGCCCTGTGGATAACAA  22
                ||||||||||||||||||||||
Sbjct  4205199  CGCACTGCCCTGTGGATAACAA  4205220
```

**Query= OriCR**

```
Length=22
>contig_2_pilon
Length=4491350

 Score = 44.1 bits (22),  Expect = 2e-06
 Identities = 22/22 (100%), Gaps = 0/22 (0%)
 Strand=Plus/Minus

Query  1        CCCTCATTCTGATCCCAGCTTA  22
                ||||||||||||||||||||||
Sbjct  4205313  CCCTCATTCTGATCCCAGCTTA  4205292
```
